# Identifying novel associations in GWAS by hierarchical Bayesian latent variable detection of differentially misclassified phenotypes

**DOI:** 10.1101/536532

**Authors:** Afrah Shafquat, Ronald G. Crystal, Jason G. Mezey

**Affiliations:** Department of Biological Statistics and Computational Biology, Cornell University, Ithaca, NY, USA; Department of Genetic Medicine, Weill Cornell Medicine, New York, NY, USA; Department of Medicine, Weill Cornell Medicine, New York, NY USA

## Abstract

Heterogeneity in definition and measurement of complex diseases in Genome-Wide Association Studies (GWAS) may lead to misdiagnoses and misclassification errors that can significantly impact discovery of disease loci. While well appreciated, almost all analyses of GWAS data consider reported disease phenotype values as is without accounting for potential misclassification. Here, we introduce Phenotype Latent variable Extraction of disease misdiagnosis (PheLEx), a GWAS analysis framework that learns and corrects misclassified phenotypes using structured genotype associations within a dataset. PheLEx consists of a hierarchical Bayesian latent variable model, where inference of differential misclassification is accomplished using filtered genotypes while implementing a full mixed model to account for population structure and genetic relatedness in study populations. Through simulations, we show that the PheLEx framework dramatically improves recovery of the correct disease state when considering realistic allele effect sizes compared to existing methodologies designed for Bayesian recovery of disease phenotypes. We also demonstrate the potential of PheLEx for extracting new candidate loci from existing GWAS data by analyzing epilepsy and bipolar disorder phenotypes available from the UK Biobank dataset, where we identify new candidate disease loci not previously reported for these datasets that have biological connections to the disease phenotypes and/or were identified in independent GWAS. In the discussion, we consider both the broader consequences and importance of careful interpretation of misclassification correction in GWAS phenotypes, as well as potential of PheLEx for re-analyzing existing GWAS data to make novel discoveries.

**Author Summary:** Prevalent misdiagnosis of diseases due to lack of understanding and/or gold-standard diagnostic measures can impact any analytics that follow. These misdiagnosis errors are especially significant in the domain of psychiatric or psychological disorders where the definition of disease and/or their diagnostic tools are always in flux or under further improvement. Here, we propose a method to extract misdiagnosis from disease and infer the correct disease phenotype. We examined the performance of this method on rigorous simulations and real disease phenotypes obtained from the UK Biobank database. We found that this method successfully recovered misdiagnosed individuals in simulations using a carefully designed hierarchical Bayesian latent variable model framework. For real disease phenotypes, epilepsy and bipolar disorder, this method not only suggested an alternate phenotype but results from this method were also used to discover new genomic loci that have been previously showed to be associated with the respective phenotypes, suggesting that this method can be further used to reanalyze large-scale genetic datasets to discover novel loci that might be ignored using traditional methodologies.

## Introduction

Identification of statistical associations between phenotypes and genotypes within Genome-wide Association Studies (GWAS) has resulted in the detection of numerous candidate genetic loci that may impact human diseases and other aspects of human physiology[1, 2]. Since the first major GWAS were published[3-6], there has been an increased realization that for many diseases and traits, it will be challenging to identify the bulk of contributing genetic loci due to the nature of genetic effects where issues include small allelic effect sizes, genetic and environmental interactions, and segregation of contributing loci for rare alleles[7, 8]. This realization has driven improved strategies for GWAS discovery such as consortium studies with large sample sizes aimed at increasing power to detect small effect size loci[9-11], sampling of understudied populations to identify loci with differential impacts due to genetics and environment[12-15], and whole-genome sequencing of individuals to assess the impact of rare alleles[16-21]. These strategies have been complemented by continued innovation in analysis methodologies applied to GWAS, including methods that detect epistatic interactions among genetic loci[22-24] and genotype by environment interactions[25-28], as well as methods aimed to extract impact of loci with rare variants[29-33]. Together, these innovations in GWAS design and methodology have led to discovery of candidate loci where the impact is particularly noticeable in diseases such as type 2 diabetes and schizophrenia where large-scale consortia studies have enabled isolation of numerous causal loci with low frequency and small effects[2, 34-37]. While these successes justify continued investment in GWAS, it is clear that continued rate of discovery of new loci for well-studied diseases and phenotypes will depend on innovative strategies that leverage underutilized aspects of GWAS.

A core aspect of GWAS that could be targeted with improved strategies is the phenotype, where there are opportunities for improved phenotype definition[38-40], measurement[41-43], and analysis[44-46]. It is well appreciated that the combination of inconsistency in methods used to diagnose disease[47, 48] and the application of imprecise measurement methodology[43] can introduce phenotyping errors that can reduce discovery potential of a GWAS[49-56]. For example, high misdiagnosis rates have been estimated for disease phenotypes such as Alzheimer’s disease and bipolar disorder which may be misdiagnosed with other forms of dementia and unipolar depression or borderline personality disorder, respectively, due to overlap of symptoms and/or lack of application of Diagnostic Systems Manual criteria[57-64]. As another example, multiple sclerosis has been frequently misdiagnosed with migraine, fibromyalgia and psychogenic disorder due to overlap of symptoms and mistakes in application of clinical and radiographic diagnostic criteria[65]. Though various strategies have been proposed to process GWAS phenotype data[46, 66-69], innovations designed to consider alternative phenotypes derived from leveraging structure of total GWAS data are particularly promising for improving candidate loci discovery in GWAS, especially when considering potential for immediate impact and implementation at minimal cost. An underexplored analysis strategy within this class of methods are those that consider misclassification of disease phenotypes[70-73], where error in disease phenotype would result in disease cases recorded as controls and vice versa[74]. Considering disease misdiagnosis rates[75, 76], there is significant potential for disease misclassification in GWAS phenotype data where even small numbers of these errors can have significant impact on GWAS statistical power and Type I errors[49, 50].

There has been surprisingly little attention paid to phenotype misclassification analysis in GWAS, where misclassification errors could be inferred and corrected by making use of genotype associations with phenotype[49, 73, 77]. The only major published methods are Bayesian approaches for recovering nondifferential misclassification (i.e., sensitivity and specificity are considered the same for cases and non-cases/controls)[49, 73] and differential misclassification (sensitivity and specificity are considered different for cases versus non-cases/controls)[77] in GWAS phenotypes. While these methods and their extensions for gene expression data have since been applied in several studies demonstrating potential benefits of misclassification analysis in identifying misdiagnosis of Alzheimer’s patients based on differential gene expression[78, 79], predicting disease subtypes in breast cancer using gene expression data[80], and finding misclassified individuals and estimating SNP effects in simulated GWAS data[49, 73, 77], a number of gaps still remain for the analysis of GWAS data. For example, most applications of existing methods are limited to the extraction of nondifferential misclassification from gene expression datasets and the only application of differential misclassification is presented in simulated GWAS data. Additionally, existing methods fail to account for inherent genetic relatedness and population structure in sampled GWAS populations, which when ignored impede GWAS discovery[81, 82]. Most importantly, these proposed methods for GWAS were only shown to perform well on GWAS datasets simulated with an artificially high number of disease-associated single nucleotide polymorphisms (SNPs) out of the total number of SNPs (i.e. only 150/1000) with genotype-specific disease-odds ratio in the range 4-10. Such simulation scenarios provide an unrealistic picture of the algorithm’s expected performance on real GWAS datasets. These previous methods also failed to directly make use of their improved phenotype classifications to discover new loci from real GWAS datasets, such that real potential for this strategy to make new discoveries from analysis of existing GWAS data has yet to be realized.

Here, we present a complete framework for Bayesian latent variable misclassification analysis to make new GWAS discoveries: Phenotype Latent Variable Extraction of disease misdiagnosis (PheLEx) (Fig 1). The core of PheLEx is a single modeling framework allowing for differential misclassification in GWAS phenotypes with an underlying full mixed model to account for population structure and genetic relatedness. When concentrating only on the problem of phenotype misclassification, we show that the PheLEx framework dramatically improved performance when analyzing simulated GWAS data that included realistic proportions of disease-associated genotypes in a genome-wide scan and genotype-specific disease-odds ratios consistent with empirical observation[83-85]. Other applications of PheLEx include exploring differential patterns between misclassified and non-misclassified cases within GWAS datasets that may point to potential causes such as misdiagnosis or disease subtypes. Furthermore, we also propose a novel strategy for applying the PheLEx framework to discover new loci within a GWAS dataset by making use of misclassification probabilities for phenotype and strategic filtering of SNPs to improve accuracy and avoid model overfitting. We demonstrate the potential of this innovative application by using PheLEx to analyze datasets for epilepsy and bipolar disorder, where we discover novel candidate loci that were not identified in the traditional analysis of these datasets. While careful interpretation of such PheLEx driven discoveries is critical, these results demonstrate potential of PheLEx for re-analyzing existing GWAS data to make novel discoveries.

**Fig 1.**
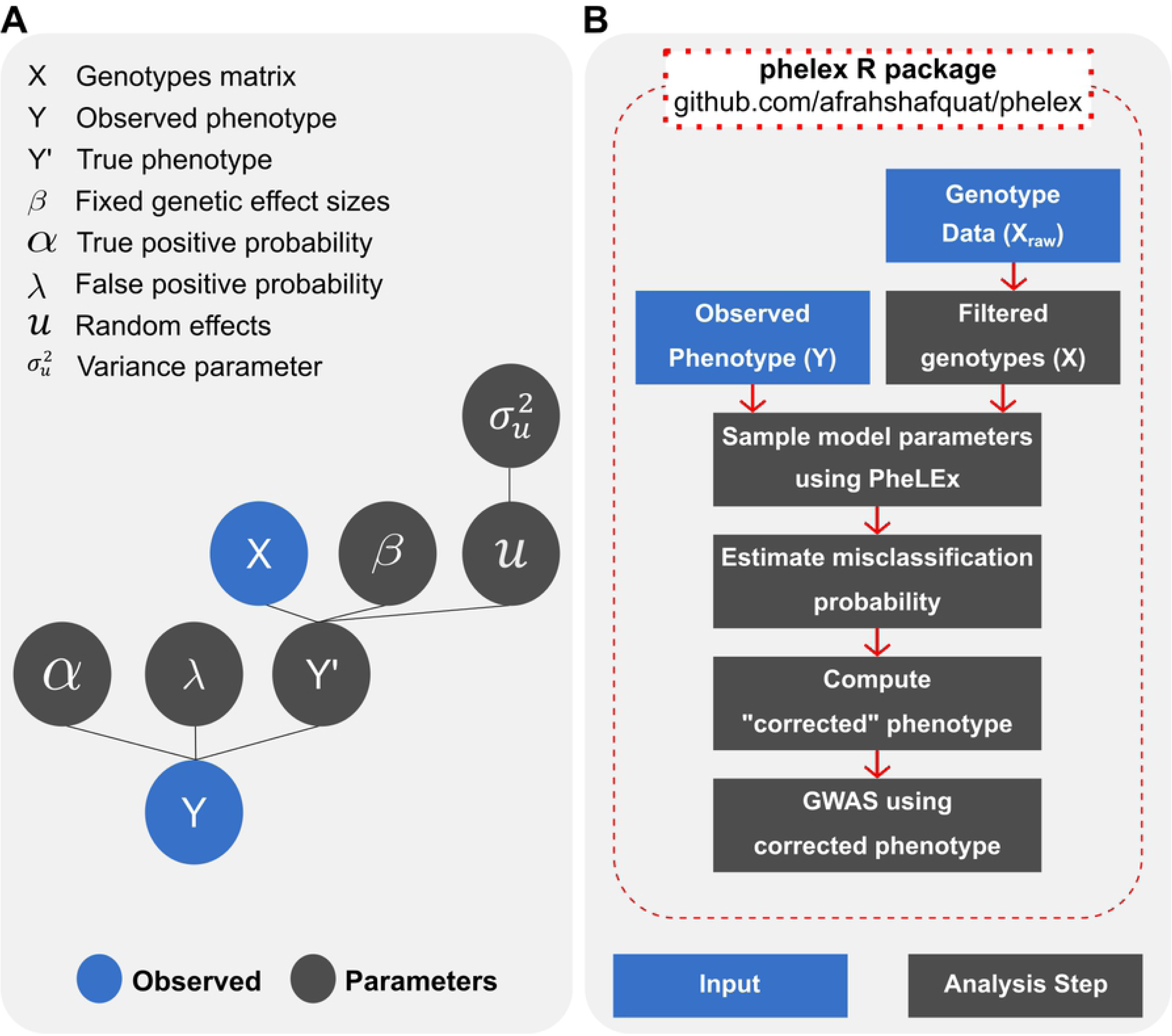
Overview of the PheLEx framework. (A) Underlying graphical model for the PheLEx method shows input parameters: genotypes matrix *X*, observed phenotype *Y*, and architecture of parameters used to infer misclassification probabilities in cases. (B) Overview of computational pipeline used to extract misclassification probabilities and produce corrected phenotypes for re-analysis using GWAS with the method PheLEx implemented in R package phelex. For detailed explanation, please refer to the main text.

## Results

PheLEx addresses limitations of existing methodologies[49, 73, 77] for Bayesian recovery of disease misclassification by introducing: (i) a more efficient Markov Chain Monte Carlo (MCMC) sampling algorithm, (ii) filtration of potentially uninformative genotypes to address issue of disproportionate ratio of disease-associated SNPs in GWAS, and (iii) accounting for genetic relatedness and population structure. Using Adaptive Metropolis-Hastings within Gibbs sampling instead of Gibbs Sampling to sample model parameters (i.e., SNP effect sizes, true-positive rate, false-positive rate and random effects) allows PheLEx more flexibility in sampling parameter estimates from landscape of posterior probabilities. As accuracy of misclassification probability under the misclassification model depends on estimated function of SNP effects and typically most SNPs in a linkage disequilibrium (LD) -pruned GWAS dataset are not associated with phenotype of interest, PheLEx filters out potentially uninformative SNPs by taking a subset of statistically significant GWAS genotypes as input, providing significant advances in terms of computational expense and misclassification accuracy. As population structure and genetic relatedness are a reality of most GWAS datasets[81, 82, 86], PheLEx accounts for these effects, which is critical for estimating accurate misclassification probabilities.

Beyond these methodology improvements for identifying misclassified phenotypes within a GWAS dataset, we also introduce a novel application of PheLEx for identifying novel GWAS associations when making use of corrected phenotypes. PheLEx presents functions that can be used to estimate misclassification probabilities to produce a corrected phenotype, which in turn can be used to perform association analysis with the genotypes matrix. The corrected phenotype provides an alternative phenotype for association, allowing for new GWAS discoveries to be made with the new phenotype. Given that PheLEx uses a subset of genotypes to estimate misclassification probabilities, any SNPs not included in this training set (and not in LD with training SNPs) that are found to be statistically significant are considered novel discoveries when analyzing the corrected phenotype.

### Simulation results

While the complete PheLEx framework includes multiple improvements over existing methodologies, to produce a fair comparison of the improved performance achieved by PheLEx compared to these methodologies[49, 73, 77], we introduced the following versions: (i) PheLEx-mh that incorporated mixed model into existing method by Rekaya et al[77] and (ii) PheLEx-m that excluded mixed model from the method PheLEx. PheLEx-mh, PheLEx-m, Rekaya (existing method) and PheLEx were applied to the simulation datasets to identify misclassified samples from increasingly misclassified simulated phenotypes (misclassification rates 5% to 40%). Differences in performance of PheLEx-m versus Rekaya and PheLEx versus PheLEx-mh showed improvement due to use of alternative MCMC algorithm (Adaptive Metropolis-Hastings within Gibbs sampling over Gibbs Sampling). Comparisons between performance of PheLEx versus PheLEx-m and PheLEx-mh versus Rekaya showed improvement when accounting for mixed effects due to population structure and genetic relatedness.

### Identification of misclassified samples in simulations

For the simulated datasets with genetic relatedness/population structure across varying degrees of misclassification (5% to 40%), PheLEx outperformed PheLEx-m, PheLEx-mh and Rekaya while the first three methods also showed superior performance to Rekaya (Figs 2A-E, S1 Fig). Area under Precision-Recall curve (AUC) values computed across simulations were also consistent on performance difference observed with PheLEx having the highest mean AUC across misclassification levels (Fig 2F), where median AUC values (listed in order of increasing misclassification) for PheLEx (median AUC: 0.552, 0.614, 0.661, 0.657 and 0.643), PheLEx-m (median AUC: 0.134, 0.239, 0.403, 0.543 and 0.600) and PheLEx-mh (median AUC: 0.467, 0.537, 0.573, 0.575 and 0.576) were higher than median AUC values for Rekaya (median AUC: 0.0676, 0.125, 0.228, 0.329 and 0.408). Performance comparisons using Receiver Operating Characteristic (ROC) curves mirrored these Precision-Recall results (S1 Fig). Additional analysis in datasets simulated without genetic relatedness/population structure also show consistent results (S1 Text).

**Fig 2.**
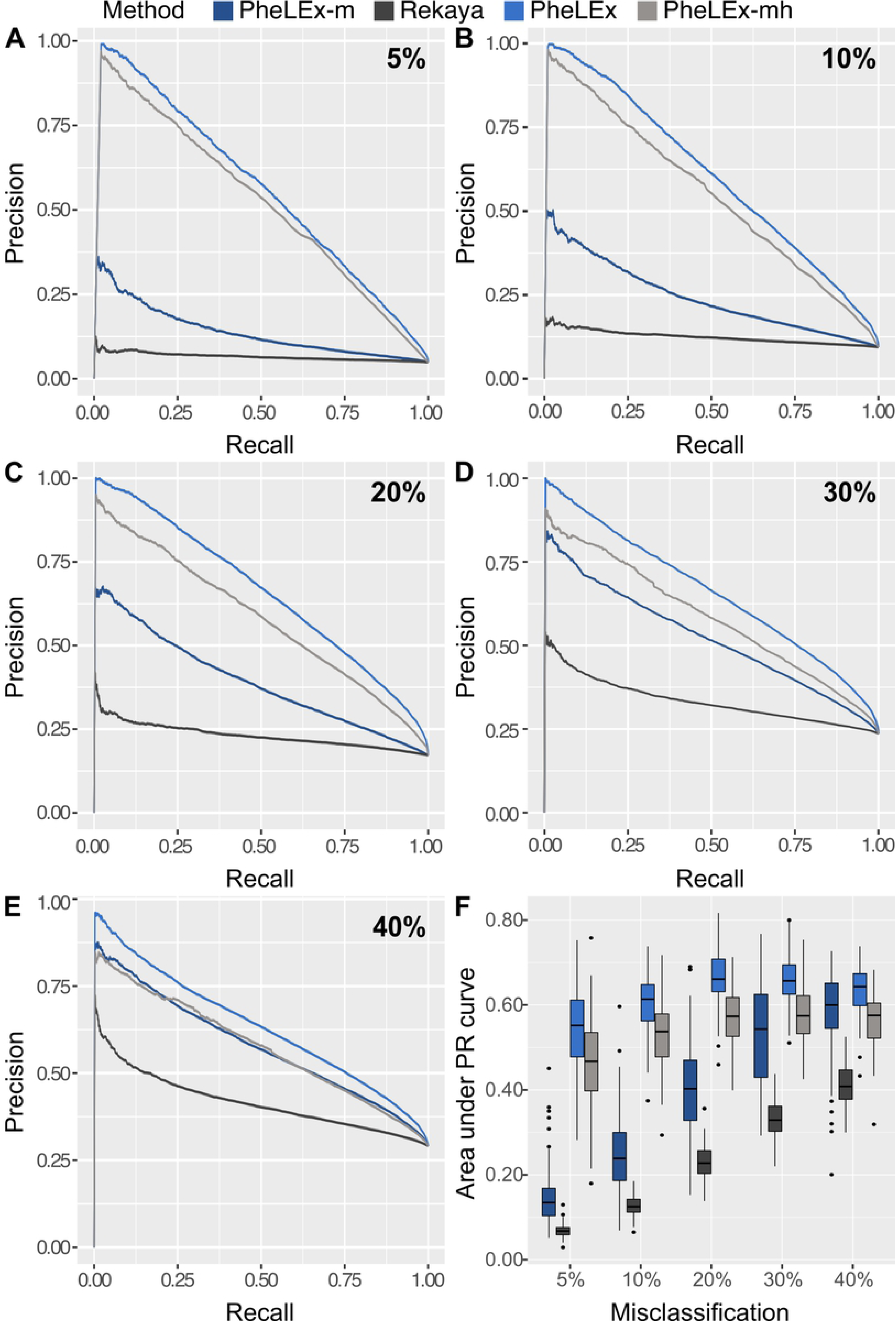
Performance comparison between PheLEx and other misclassification extraction methods for simulated GWAS datasets. (A-E) Mean precision (y-axis) over recall (x-axis) curves over 100 simulated datasets are shown for increasing misclassification rates of 5%, 10%, 20%, 30% and 40%. (F) Boxplots for area under Precision-Recall (PR) curves (AUC) (y-axis) are shown for 100 simulated datasets across increasing misclassification rates (x-axis). Misclassification extraction methods are indicated with different colors: PheLEx-m (dark blue), Rekaya (dark gray), PheLEx (light blue), and PheLEx-mh (light gray).

Accounting for population structure and genetic relatedness resulted in greatest increase in precision over recall as shown by performance of PheLEx-mh versus Rekaya and PheLEx versus PheLEx-m. Though use of Adaptive Metropolis-Hastings improved performance, this boost was not on the same scale as observed by introducing the mixed model. Though overall performance improved with increasing rates of misclassification, improvement in performance plateaued around 30%-40% rates of misclassification. This plateau in performance can be explained by reduction in total number of informative SNPs passing filtration criteria due to growing misclassification rates, thus affecting estimated SNP effects and consequently, accuracy of misclassification probabilities inferred.

Improvement in performance was accompanied by a boost in speed as a result of two important components of the PheLEx framework i.e., filtration of SNPs and use of Adaptive Metropolis-Hastings within Gibbs sampling. Using results from applying misclassification extraction methods (i.e., PheLEx, PheLEx-mh, Rekaya and PheLEx-m) to simulation datasets with genetic relatedness/population structure, we observed running time for each method (S2 Fig). Across all simulations, PheLEx (median time: 31.2 minutes) was more efficient than PheLEx-mh (median time: 601 minutes) while PheLEx-m (median time: 6.34 minutes) was faster than Rekaya (median time: 41.6 minutes). Though accounting for mixed effects adds to running time as shown by comparison between PheLEx-m and PheLEx, PheLEx’s speed was still comparable to that of Rekaya underlining efficiency of our MCMC algorithm of choice.

### GWAS discovery via correction by PheLEx in simulated datasets

To explore the impact of identifying misclassified samples on association analysis, we computed corrected phenotypes using misclassification probabilities obtained from PheLEx for simulated data. Corrected phenotypes were produced by switching cases with high misclassification probabilities (determined using thresholds defined in the methods section) to controls using PheLEx. We performed GWAS on simulated true phenotypes (no misclassification), misclassified phenotypes and PheLEx corrected phenotypes, and quantified the ratio of sum of statistically significant disease-associated SNPs (according to Bonferroni-corrected p-value threshold) for misclassified phenotypes and corrected phenotypes against sum of statistically significant disease-associated SNPs for the simulated true phenotypes (no misclassification).

As expected, with increasing misclassification, this ratio decreased for misclassified and corrected phenotypes signifying disease-associated SNPs that lose statistical significance in GWAS (Fig 3A). However, the ratio (listed in order of increasing misclassification) for PheLEx corrected phenotypes (mean ratio: 0.951, 0.894, 0.776, 0.629 and 0.469) improved upon the ratio for misclassified phenotypes (mean ratio: 0.951, 0.891, 0.765, 0.610 and 0.454) at each misclassification level with increased improvements at higher misclassification rates (20-40%). Smaller fractions of disease-associated SNPs were recovered at low misclassification rates as compared to higher misclassification rates. At low misclassification rates, lower precision of switching cases would entail loss of true cases (switched to controls by PheLEx) along with misclassified cases in corrected phenotype, reducing PheLEx’s ability to recover true disease-associated SNPs.

**Fig 3.**
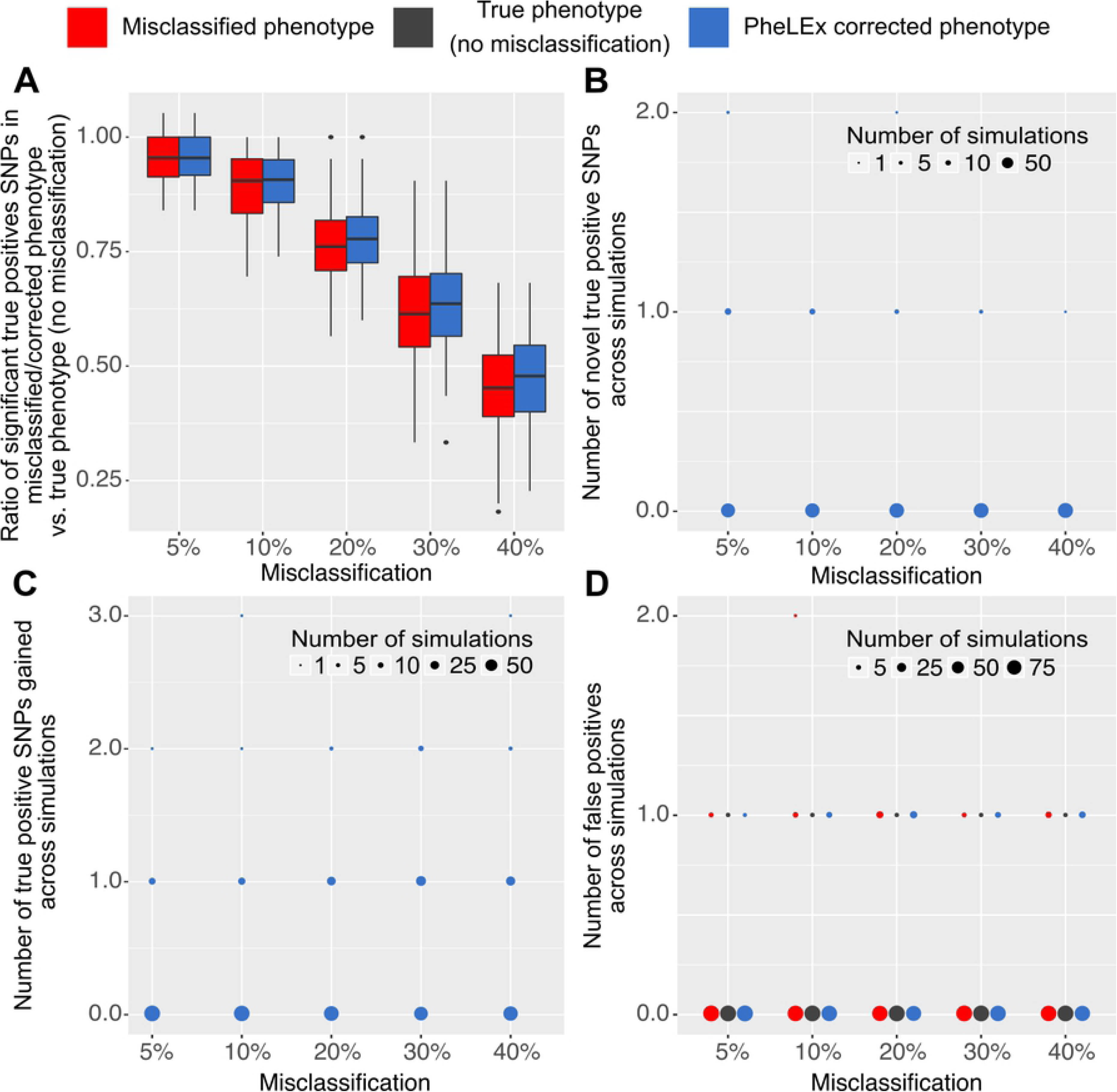
GWAS results using PheLEx corrected phenotypes for simulated datasets. (A) Boxplots of the ratio of sum of significant disease-associated SNPs for the misclassified phenotype versus simulated true phenotype (no misclassification), and ratio of sum of significant disease-associated SNPs for the PheLEx corrected phenotype versus in simulated true phenotype (no misclassification) (y-axis) is shown for increasing misclassification rates (x-axis). (B) Number of novel disease-associated SNPs (y-axis) is shown across simulations for increasing misclassification rates (x-axis). (C) Number of true positive SNPs gained (y-axis) across simulations is shown against misclassification rates (x-axis). (D) Number of false positives (y-axis) are shown across simulations for increasing misclassification rates (x-axis). Size of points is proportional to the number of simulations the values of false positive/true positive SNPs gained/novel SNPs are found within and color of points indicates GWAS results for misclassified phenotype (red), true phenotype (no misclassification) (dark gray) and PheLEx corrected phenotype (blue).

Importantly, we were also able to identify novel SNPs defined as disease-associated SNPs that were not statistically significant for either the simulated true phenotypes (no misclassification) or misclassified phenotypes but were statistically significant for PheLEx corrected phenotypes (Figs 3B and 3C), where these novel discoveries were not accompanied by recovery of large numbers of false positives (Fig 3D). Given the modest false positives comparable to those already found in misclassified phenotypes and improvement in discovery of statistically significant disease-associated SNPs, these results indicate PheLEx is a viable approach for making new discoveries in existing GWAS datasets.

### Finding novel associations in real GWAS datasets

PheLEx was applied to UK Biobank GWAS datasets for epilepsy and bipolar disorder to extract misclassification probabilities for the phenotype cases. Using a threshold, misclassified cases were identified for each phenotype and their respective phenotypes were switched from case to control, resulting in corrected phenotypes. Using these corrected phenotypes, we performed association analysis to investigate the genetic associations with the new phenotype. In both analyses, we observed improvement in statistical power of association analysis and identification of novel loci in GWAS.

### Epilepsy

We analyzed UK Biobank dataset for epilepsy phenotype using PheLEx to identify *n* = 597 individuals whose phenotypes might be misclassified. These cases were identified as potentially misclassified and their phenotype switched from cases to controls to compute a “corrected” epilepsy phenotype. GWAS was performed on original epilepsy phenotype of dataset and corrected epilepsy phenotype was produced by PheLEx to compare results (Fig 4). Results of the original analysis were similar to that produced previously for this dataset[87] with no statistically significant SNPs according to Bonferroni-corrected p-value threshold. However, after correction of phenotype an overall improvement in statistical significance of SNPs can be observed. Apart from training SNPs, SNPs not used in training also gained statistical significance at a Benjamini-Hochberg adjusted p-value < 0.1. By computing the r^2^ measure of linkage disequilibrium (LD) amongst these candidate SNPs and training SNPs (Fig 5), we were able to extract three interesting candidate SNPs not in LD with training SNPs that gained statistical significance (Table 1). Even though most SNPs underwent relatively small changes in their p-values (in either direction), the three candidate SNPs experienced a significant boost from their original p-values indicating the potential of PheLEx to discover new loci. These candidates included SNP in gene *PRUNE2* (MIM: 610691) that has been significantly associated in an independent GWAS for hippocampal atrophy[88] linked to epilepsy [89, 90], and has also been associated with other neurological phenotypes in GWAS for mental retardation and Alzheimer’s disease[88, 91]. The closest genes near two other candidate SNPs include *ERCC6* which is an epilepsy-related gene[92] and *OR2S2*, which has not been previously linked to epilepsy. When looking at SNPs in LD with candidate SNP rs12379440 (SNP in *OR2S2*), these include SNPs in genes *AK3* (MIM: 609290), *DIRAS2* (MIM: 607863), *NTRK2* (MIM: 600456), *MAMDC2* (MIM: 612879) and *OSTF1* (MIM: 610180). At least one or more of these genes have associations with seizure phenotypes[93-95], attention deficit hyperactivity disorder (ADHD)[96]and temporal lobe epilepsy[97].

**Table 1.** Novel candidate SNPs identified for UK Biobank epilepsy phenotype.

| SNP ID | Location<br>(chromosome:<br>position) | Gene | SNP p-value in<br>original phenotype | SNP p-value in<br>corrected phenotype |
| --- | --- | --- | --- | --- |
| rs12379440 | 9:35963328 | *OR2S2*<sup>a</sup> | 2.27e-4 | 4.87e-6 |
| rs17778913 | 9:79274466 | *PRUNE2* | 7.89e-4 | 6.73e-7 |
| rs34535659 | 10:50751024 | *ERCC6-*<br>*PGBD3*<sup>a</sup> | 1.11e-4 | 4.82e-6 |

<sup>a</sup> Nearest HUGO Gene Nomenclature Committee (HGNC) gene is reported as SNP wasn't found within a HGNC gene. All annotations were performed using web-resource snpSNAP[98]

**Fig 4.**
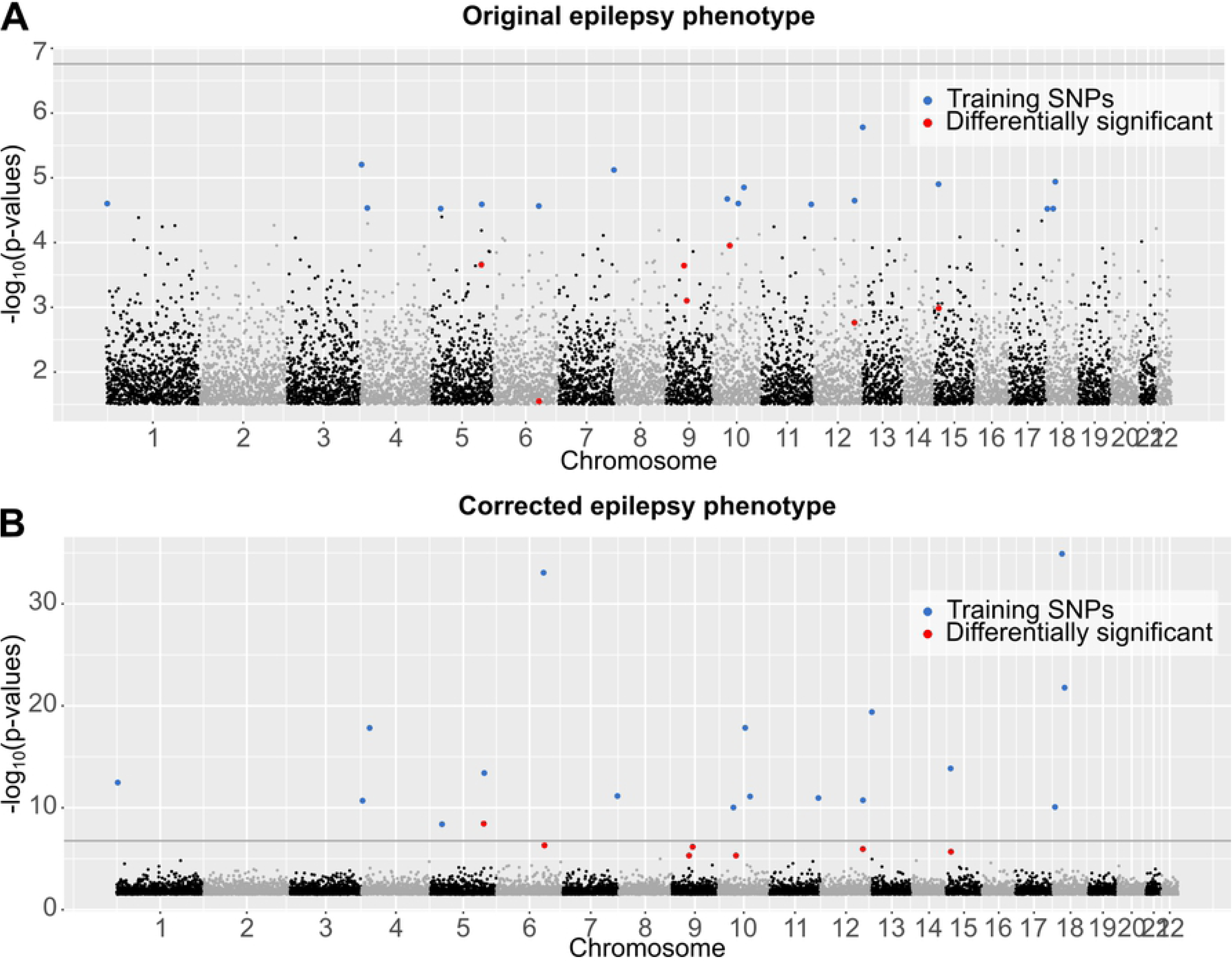
Association analysis of epilepsy dataset using PheLEx. Manhattan plots (x-axis: SNP genomic position, y-axis: −log_10_ p-values of association test > 1.52) of (A) GWAS results for original epilepsy phenotype with Bonferroni-corrected p-value threshold shown as dark gray line, (B) GWAS results for PheLEx corrected phenotype (where PheLEx-identified cases are switched to controls). Training SNPs used as input for PheLEx are marked in blue whereas differentially significant SNPs are marked in red. Differentially significant SNPs are defined as SNPs not included in training PheLEx that are statistically significant using Benjamini-Hochberg procedure (adjusted p-value < 0.1) in the PheLEx corrected epilepsy phenotype and not in original epilepsy phenotype.

**Fig 5.**
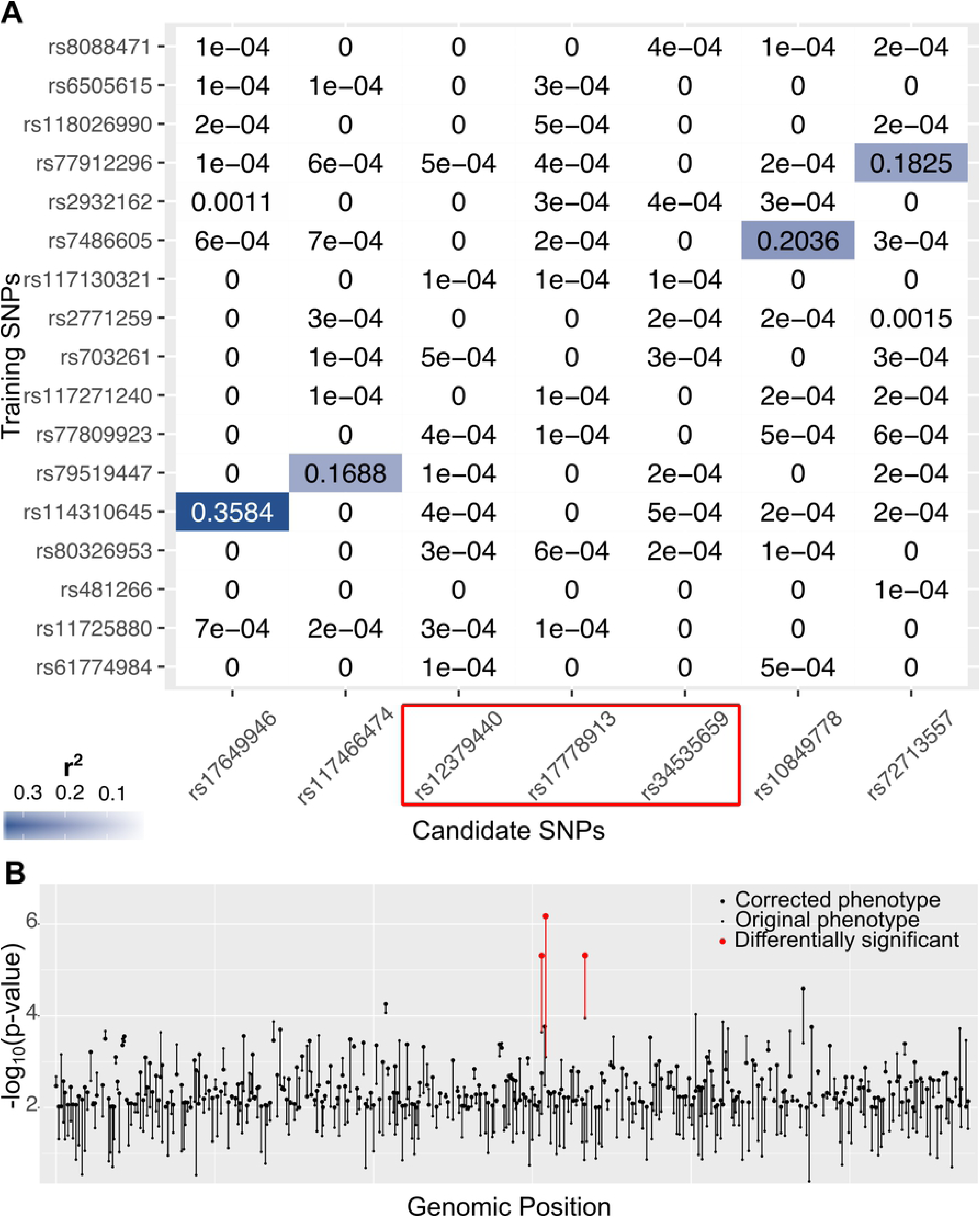
Candidate SNPs for epilepsy phenotype. (A) Heatmap showing r^2^ measure of linkage disequilibrium (LD) computed between the seven differentially significant SNPs (columns) identified for corrected epilepsy phenotype and training SNPs used as input for PheLEx (rows). Candidate SNPs with r^2^< 0.1 are marked within a red box indicating three differentially significant SNPs in corrected epilepsy phenotype that are not in LD with training SNPs. (B) Log transformed p-values (y-axis) are reported for a subset of SNPs not in LD with training SNPs plotted in genomic position (x-axis). The small dot for each SNP denotes −log_10_(p-value) in original epilepsy phenotype and large dot denotes −log_10_(p-value) in the PheLEx corrected epilepsy phenotype. Differentially significant SNPs (adjusted p-value < 0.1) in the PheLEx corrected epilepsy phenotype compared to original epilepsy phenotype are represented in red.

### Bipolar disorder

We analyzed UK Biobank data for bipolar disorder phenotype using PheLEx and identified *n* = 179 cases that might be potentially misclassified. Corrected phenotype was produced where cases identified using PheLEx as misclassified were changed to controls. Although, GWAS results with original bipolar disorder phenotype failed to produce any statistically significant SNPs (consistent with previous analysis[87]), results from the corrected phenotype identified candidate loci with statistical significance at a Benjamini-Hochberg adjusted p-value < 0.1 (Fig 6). All identified candidate loci were not in LD with training SNPs and experienced substantial improvement in their statistical significance from original phenotype to corrected phenotype (Fig 7). Table 2 lists details for these candidate SNPs including the genes they were located within or nearest. Though one of candidates SNPs was found within gene *NR3C1* (MIM: 138040) which is directly linked to seizure phenotypes[99], the other candidate SNPs were located in genes *RNF168* (MIM:612688) and *FBP1* (MIM: 611570), which have not been associated directly with bipolar disorder. However, a candidate SNP in *RNF168* is in LD with SNPs in *ATP13A4* (MIM: 609556) and *KCNH8* (MIM: 608260), where both genes have been linked to schizophrenia[100-102]. Similarly, SNPs in *FBP1* are in LD with candidate SNP rs7853477 which has been linked to psychosis and schizophrenia[103, 104].

**Table 2.** Novel candidate SNPs identified for UK Biobank bipolar disorder phenotype.

| SNP ID | Location<br>(chromosome:<br>position) | Gene | SNP p-value in<br>original phenotype | SNP p-value in<br>corrected phenotype |
| --- | --- | --- | --- | --- |
| rs116629271 | 3:183453024 | *SNORA63E* | 1.69e-4 | 2.87e-5 |
| rs13090871 | 3:196202326 | *RNF168* | 1.12e-4 | 5.12e-7 |
| rs72801072 | 5:142720593 | *NR3C1* | 1.65e-4 | 2.15e-5 |
| rs117958051 | 8:49696665 | *EFCAB1*<sup>a</sup> | 5.80e-5 | 3.48e-6 |
| rs7853477 | 9:91530248 | *PCNPP2*<sup>a</sup> | 9.80e-4 | 1.66e-5 |

<sup>a</sup> Nearest HUGO Gene Nomenclature Committee (HGNC) gene is reported as SNP wasn't found within a HGNC gene. All annotations were performed using web-resource snpSNAP[98]

**Fig 6.**
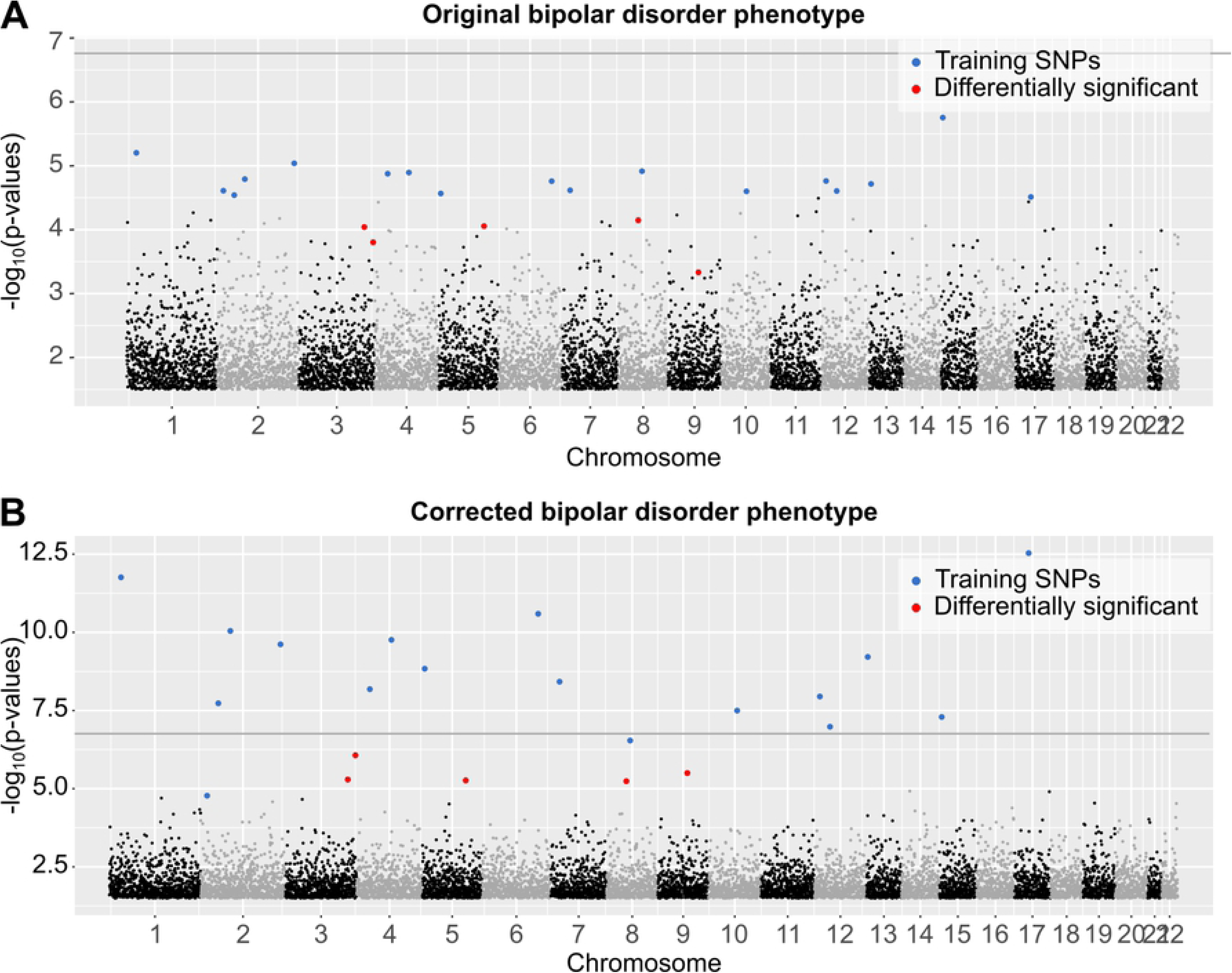
Association analysis of bipolar disorder dataset using PheLEx. Manhattan plots (x-axis: SNP genomic position, y-axis: −log_10_ p-values of association test > 1.52) of (A) GWAS results for original bipolar disorder phenotype with Bonferroni-corrected p-value threshold shown as dark gray line, (B) GWAS results for PheLEx corrected bipolar disorder phenotype (where PheLEx-identified cases are switched to controls). Training SNPs used as input for PheLEx are marked in blue whereas differentially significant SNPs are marked in red. Differentially significant SNPs are defined as SNPs not included in training PheLEx that are statistically significant using Benjamini-Hochberg procedure (adjusted p-value < 0.1) in the PheLEx corrected bipolar disorder phenotype and not in original bipolar disorder phenotype.

**Fig 7.**
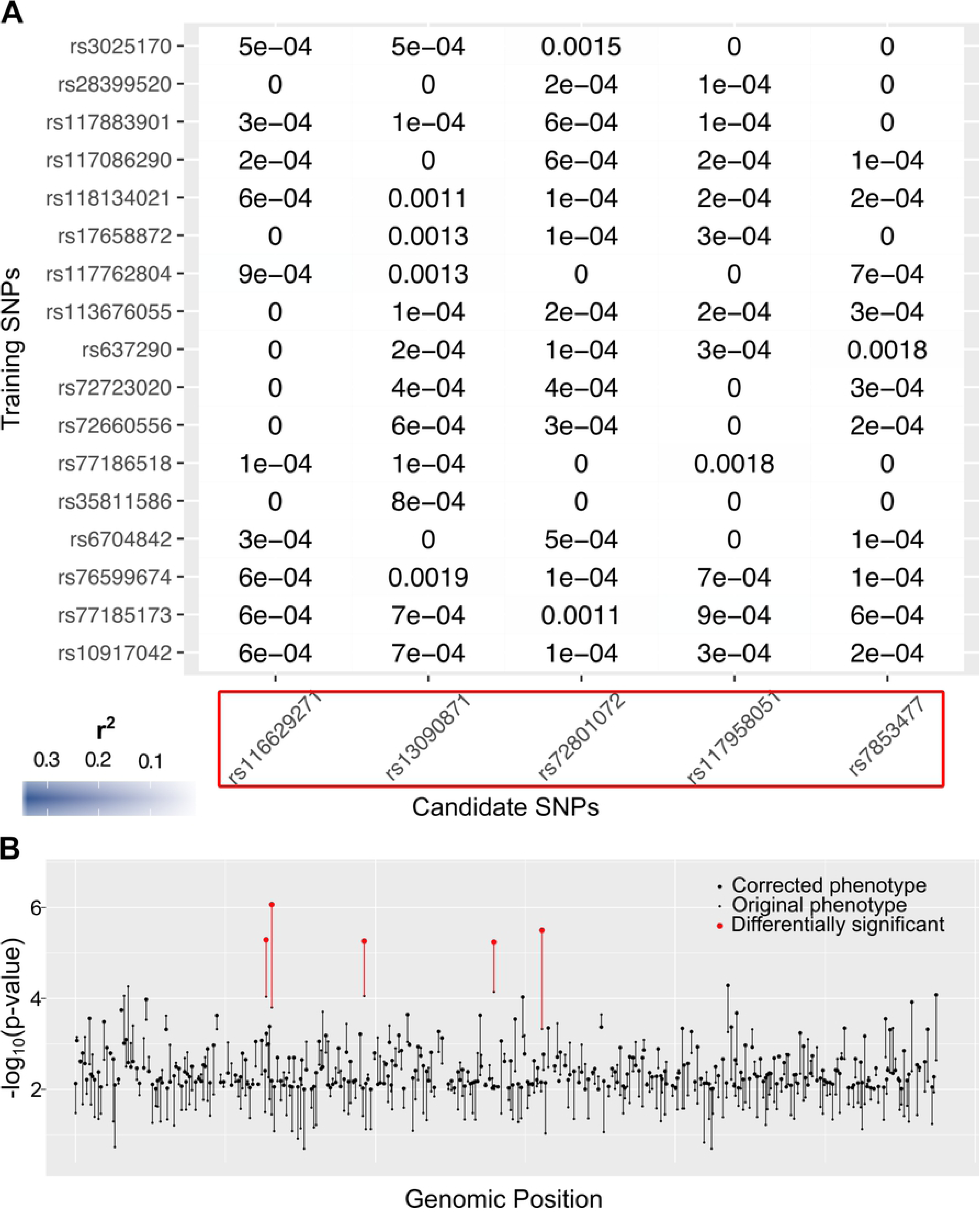
Candidate SNPs for bipolar disorder phenotype. (A) Heatmap showing r^2^ measure of linkage disequilibrium (LD) computed between five differentially significant SNPs (columns) identified for corrected bipolar disorder phenotype and training SNPs used as input for PheLEx (rows). Candidate SNPs with r^2^ < 0.1 are marked within a red box indicating five differentially significant SNPs in corrected bipolar disorder phenotype that are not in LD with training SNPs. (B) Log transformed p-values (y-axis) for a subset of SNPs not in LD with training SNPs plotted in genomic position (x-axis). The small dot for each SNP denotes −log_10_ (p-value) in original bipolar disorder phenotype and large dot denotes −log_10_ (p-value) in the PheLEx corrected bipolar disorder phenotype. Differentially significant SNPs (adjusted p-value < 0.1) in the PheLEx corrected bipolar disorder phenotype compared to original bipolar disorder phenotype are represented in red.

## Material and Methods

### Simulation datasets

Genotypes were simulated using simulateGenotypes function from R package PhenotypeSimulator[105] for 2,000 samples and 10,000 independent SNPs. Minor allele frequency (MAF) for simulated SNPs was sampled from multinomial distribution with means 0.1, 0.2 and 0.4 (default parameters for simulateGenotypes function). One hundred true disease phenotypes (*Y’*) were simulated using the following relationship:

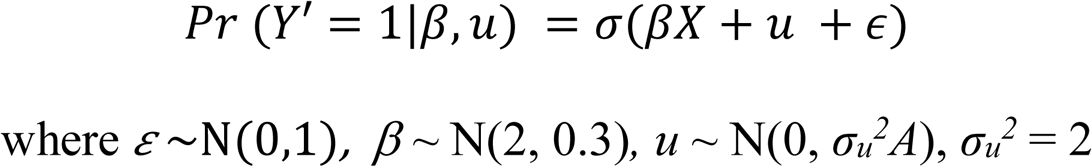

Here, *σ* is a probit link function, *β* are fixed effect sizes of disease-associated SNPs *X, u* is a simulated random effects vector, and *ε* represents noise. *A* is a square genetic relatedness matrix (*n* = 2,000) computed using getKinship function from R package PhenotypeSimulator. Random effects vector *u* was simulated from multivariate normal distribution using function mvrnorm from R package MASS with variance parameter *s*_*u*_^*2*^, relatedness matrix *A* and mean as zero vector. The same random effects vector was used to simulate all 100 true disease phenotypes *Y’*. Thirty SNPs were randomly selected to be disease-associated SNPs *X* for all true phenotypes *Y’*. Fixed effects *β* for *X* were sampled for each disease phenotype separately from a normal distribution with mean and variance parameter values stated above. The genetic model with 30 disease-associated SNPs out of total 10,000 preserves the characteristic sparsity of “true signal” in GWAS datasets, while keeping the dataset size manageable for rigorous simulations. Number of samples, number of SNPs, number of disease-associated SNPs, and effect sizes were set in accordance with precedence in literature[106-108], where in particular, effect sizes were selected to ensure genotype-specific disease odds ratio remained realistic (in the range: 1-3)[83-85] (S3 Fig). For each simulated true disease phenotype *Y’*, differential misclassification was introduced at varying degrees by switching a fraction of randomly selected controls to cases. The fraction of controls switched to cases varied from 5%, 10%, 20%, 30% and 40% representing increasing rates of misclassification in “observed phenotype” denoted as *Y*. Resulting datasets consisted of 100 datasets for each misclassification rate simulated with mixed effects of genetic relatedness/population structure. We note that this configuration of simulated GWAS datasets is in stark contrast to simulated data analyzed previously[49, 77], where 150 out of 1000 simulated SNPs were associated with true disease phenotype with maximum genotype-specific disease odds-ratio in range 4-10 in each dataset. Additionally, we simulated datasets without genetic relatedness/population structure to directly compare method performance in a less realistic case (S1 Text).

### Case studies datasets

Phenotype and genotype data for epilepsy and bipolar disorder were obtained from UK Biobank[109]. Data preprocessing steps used were similar to those adapted in previous analysis for UK Biobank datasets where any differences made no significant impact on results obtained from previous analyses of these GWAS data[87]. For a previous analysis of UK Biobank phenotypes, filtration steps included removing genotypes based on genotyping missingness rate > 2% across samples, minor allele frequency (MAF) < 10^-4^ and departure from Hardy-Weinberg Equilibrium p < 10^-50^, while samples were removed based on missingness rate > 5% across variants, inconsistency between self-reported gender and genetic sex inferred, and non-British white ancestry. For our analysis of both phenotypes, we filtered UK Biobank dataset using steps described above with the exception of MAF threshold which was replaced with a more conservative threshold of 10^-3^. Additionally, all genotypes in linkage disequilibrium (LD) were pruned from the dataset using PLINK[23] with specific parameters (flag: --indep-pairwise, window size: 50kb, step size: 5, r^2^ threshold: 0.20). This resulted in 287,425 SNPs in the UK Biobank dataset which was then divided into datasets for epilepsy and bipolar disorder phenotypes based on diagnosis record. Cases were selected based on diagnosis records where any individual containing diagnosis code for phenotype was labeled a case (bipolar disorder code: 1291 and epilepsy code: 1264). For epilepsy, 3,620 cases were identified and an equal number of controls (*n* = 3,620) were randomly selected from a pool of individuals who did not have epilepsy as their diagnosis in the UK Biobank dataset. For bipolar disorder, 1,177 cases were identified and twice the number of controls (*n* = 2,354) were randomly selected from a pool of individuals who did not have bipolar disorder as their diagnosis in the UK Biobank dataset. Distribution of phenotypes and sex within these datasets is described in S1 Table.

### Misclassification analysis of simulated and cases studies datasets

For application of misclassification extraction methods (detailed below) on simulation datasets, we filtered out potentially uninformative SNPs using their respective p-values from standard GWAS analysis using Bonferroni-corrected p-value threshold as cut-off. For all methods, resulting input genotypes matrix (training SNPs) contained SNPs that passed Bonferroni corrected genome-wide threshold for the dataset. Additionally, only PheLEx was applied to epilepsy and bipolar disorder datasets to identify potentially misclassified samples. For both real phenotypes, association analysis failed to produce any statistically significant SNPs using Bonferroni-corrected threshold. We selected a threshold (-log_10_(p-value) > 4.5) as a heuristic to filter training SNPs used as input for PheLEx. Input for misclassification extraction methods accounting for mixed effects also included genetic similarity matrix computed for simulations and real datasets using R function getKinship on SNPs with MAF > 5%[105].

### Misclassification model

In absence of misclassification in phenotype, relationship between genotypes matrix *X* (composed of *j* SNPs) and observed phenotype *Y* (for *n* individuals) can be stated as,

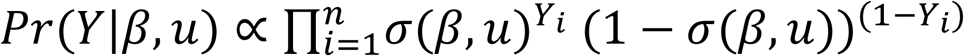

where

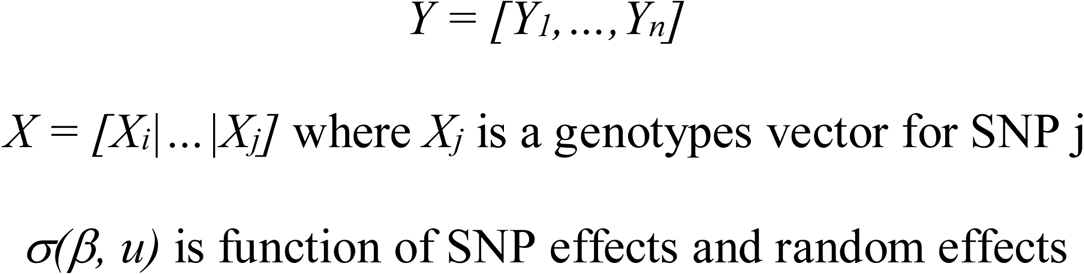

In presence of misclassification, the relationship between *X* and *Y* can be modeled using hierarchical Bayesian latent variable model (Fig 1A). Here *X* and *Y* are intermediated through (i) latent variable representing true phenotype *Y’*, (ii) false-positive rate in phenotype (*λ*) representing rate of true controls recorded as cases, and (iii) true-positive rate in phenotype (*α*) representing rate of true cases recorded as cases. Hence, the relationship between genotypes *X* and true phenotype *Y’* can be stated as:

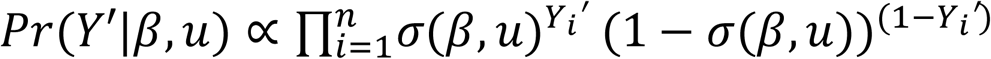

Resulting likelihood of observing the data (*X* and *Y*) given unknown parameters is:

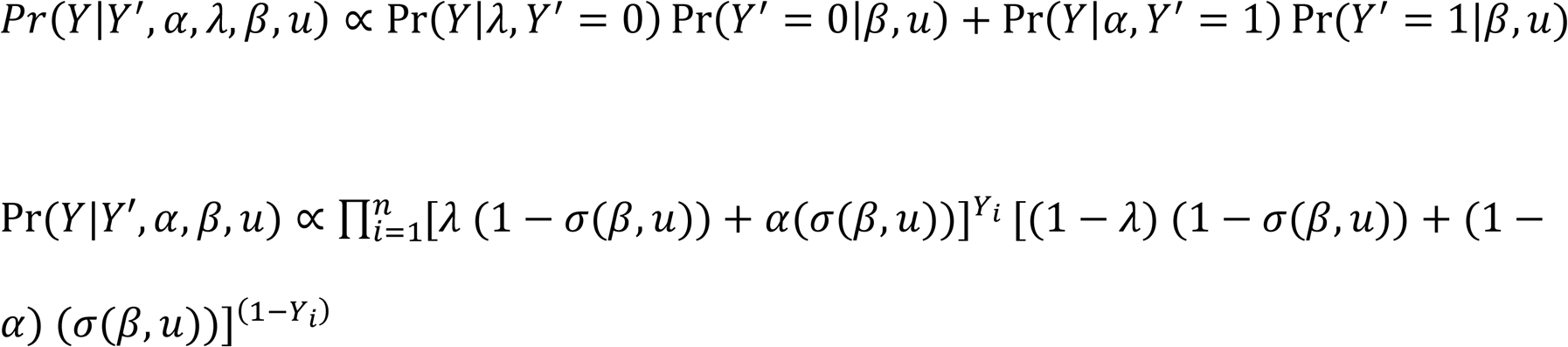

Hence, posterior probability of the model is:

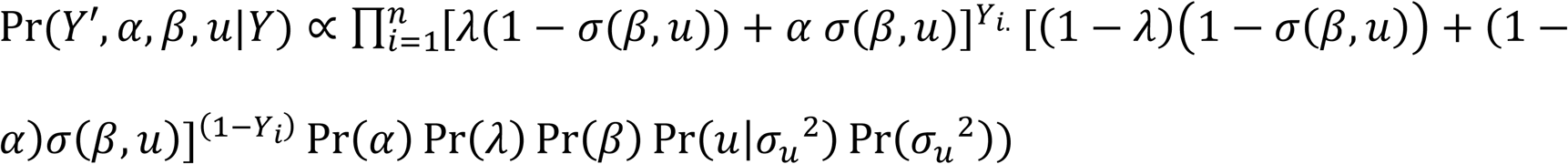

### PheLEx Algorithm

PheLEx follows a computational framework detailed in Fig 1B. PheLEx takes a subset of GWAS genotypes dataset *X* as input (training SNPs) containing SNPs based on the statistical significance of association with phenotype of interest. Other inputs for PheLEx include observed phenotype *Y* and genetic relatedness matrix *A*. Given these inputs, PheLEx employs Adaptive Metropolis-Hastings within Gibbs Sampling to estimate model parameters. Parameters (*α, λ* and *β*) are estimated in Adaptive Metropolis-Hastings step using the following steps:

1. Initialize random starting values for parameters *α, λ* and *β* using respective distributions to sample starting values. Set *u* as a zero vector and *σ*_*u*_^*2*^= 0.1
2. Define proposal
  a. Sample values for *α* and *λ* using truncated normal distribution
  b. Sample values for *β* using normal distribution
3. Calculate posterior probabilities from current parameter values and proposed parameter values
  a. Compute *σ*(β,u) for current parameter values and proposed parameter values
  b. Compute posterior for current values and proposed parameter values using

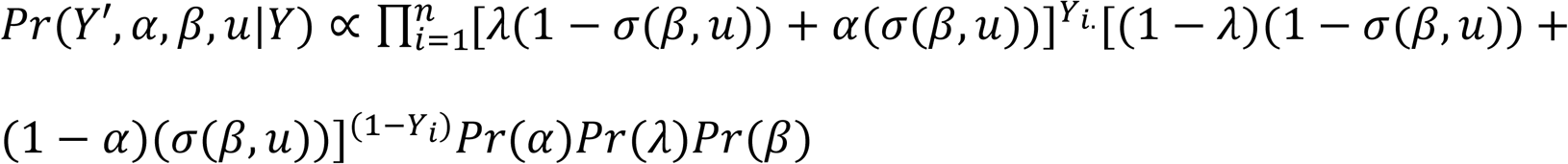
  c. Update values for parameters with proposed parameter values with probability 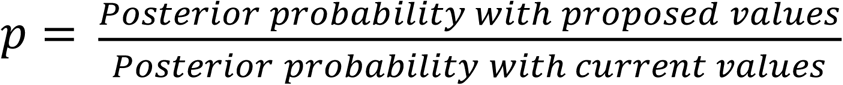

Parameters *σ* _*u*_^*2*^ and *u* are estimated in the Gibbs step using conditional probability distributions for each parameter as defined in previous literature[110-112].

1. Given *l*_*i*_ = *X*_*i*_+ *u*_*i*_
2. 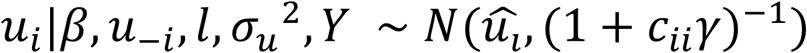 where,
  a. 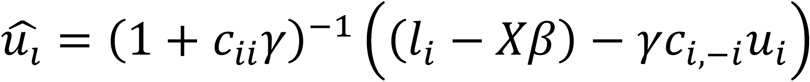
  b. *c*_*i,i*_ = *ith diagonal element of A^−1^*
  c. *c*_*i*, −*i*_ = *row i of A with element i removed*
  d. *u*_-*i*_ = *vector u with element i removed*
  e. 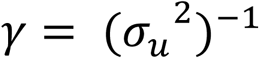
3. 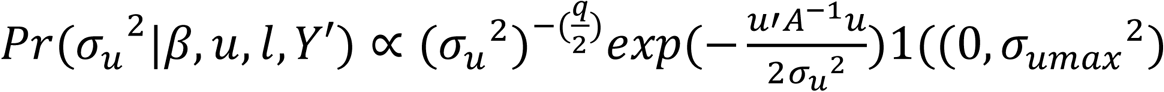
  a. *σ*_*umax*_^2^ = 100
4. Estimate misclassified phenotypes using
  a. Misclassification in cases ∼ Binomial (*n*_*1*_, Pr(*Y’*=0|*Y*=1, *α, λ, β, u*))
    i. 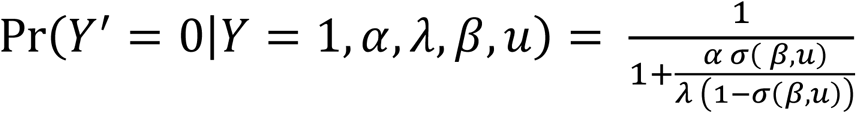
    ii. *n*_1_ = *total number of cases*
  b. Misclassification in controls ∼ Binomial (*n*_*2*_, Pr(*Y’*=1|*Y*=0, *α, λ, β, u*)))
    i. 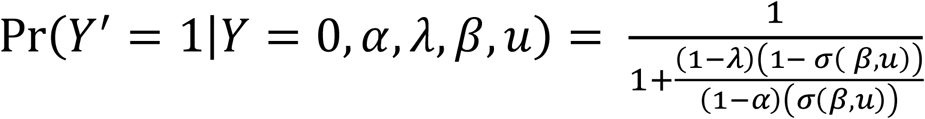
    ii. *n*_2_ = *total number of controls*

For simulations and real data, prior probabilities on parameters are defined as Pr(*α*) ∼ Beta(10, 1), Pr(*λ*) ∼ Beta(1, 1), Pr(*β*) ∼ N(1, 5) with flat prior on variance parameter *σ*_*u*_^*2*^ as suggested by[111, 112]. Variance for jumping distribution of effect sizes was adjusted across iterations to maintain acceptance ratio for Markov Chain Monte Carlo (MCMC) chains around 0.2 using established methods defined previously[113]. Acceptance rate of 0.2 was used[114] and the algorithm was run on each dataset for 100,000 iterations with a burn-in of 20,000 iterations. At each iteration, estimates for each parameter (*α, λ, β* and *u*) were used to calculate misclassification probability for each sample, which in turn was used to compute whether each sample was misclassified.

Using a defined threshold on estimated misclassification probabilities, potentially misclassified cases were identified for simulations and real datasets. Precision, recall and other operating characteristics were calculated for misclassified cases identified by PheLEx in simulations and compared to performance of other algorithms described below. Misclassified cases found in simulations and real datasets were further used to create corrected phenotypes by switching phenotype of misclassified cases from case to control. Association analysis was performed with corrected phenotypes from simulations and used to estimate impact on GWAS discovery. For real datasets, association analysis was performed with corrected phenotypes where SNPs differentially significant in analysis with corrected phenotype versus original disease phenotype were investigated for biological significance as described below.

### Other misclassification algorithms

In addition to PheLEx, we designed two additional algorithms to provide a fair comparison between PheLEx and existing methods proposed by Rekaya[49, 73, 77]. PheLEx-m (PheLEx - /minus mixed model) was designed to exclude the mixed model component of PheLEx. Comparison between PheLEx-m and Rekaya is used to determine improvement due to use of an Adaptive Metropolis-Hastings within Gibbs (used in PheLEx) over Gibbs Sampling algorithm (used in Rekaya). PheLEx-mh (PheLEx -/minus Metropolis-Hastings) was designed to replace the MCMC algorithm in PheLEx with Gibbs Sampling (as used in Rekaya), however, with the mixed model present as implemented in PheLEx. Comparison between Rekaya and PheLEx-mh provide insight into improvement due to accounting for mixed effects due to population structure and genetic relatedness as suggested by the PheLEx framework. Implementation steps for PheLEx-m, PheLEx-mh and Rekaya’s method[77] are described in S2 Text.

### Genome-wide association analysis

The following procedure was used to produce inputs for analysis of simulations and real case studies. Standard association analysis was applied using a low-rank linear mixed model as implemented in R package lrgpr[115], where for simulations no additional covariates were used in association analysis whereas for epilepsy and bipolar disorder datasets, GWAS was performed using sex (Male/Female) as an additional fixed covariate in the association model. The function used for association analysis was lrgprApply, which uses cross-validation and model selection criteria to estimate a low-rank genetic similarity matrix to be used in association analysis[115]. This analysis was similar to those performed previously for UK Biobank phenotypes[87], however, some covariates (e.g. age, batch) previously included[87] were removed from our analysis based on little improvement to quality of Quantile-Quantile plots and GWAS results (S4 Fig).

### Identifying misclassified samples in phenotype

In simulations, each algorithm (PheLEx, PheLEx-m, PheLEx-mh and Rekaya) produced a matrix of misclassification indicators using estimated misclassification probability for cases and controls computed at each iteration. The rows in the matrix represented samples and columns represented each iteration, where value 1 for any sample at any iteration indicated sample is misclassified (or 0 otherwise) according to parameter estimates at that iteration. Average misclassification probability for each sample was computed by dividing sums of each row by total iterations. Misclassified individuals were identified by using a defined threshold as cut-off on average misclassification probability (*p*) computed for each individual where if *p* is greater than threshold, the individual is determined to be misclassified. For simulations, threshold was defined as below as defined previously [77]:

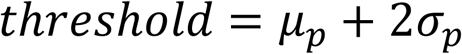

where *p* = vector of average misclassification probabilities of all cases, *μ*_*p*_ = mean of average misclassification probabilities for all cases, and *σ*_*p*_= standard deviation of average misclassification probabilities for all cases. Corrected phenotype in each simulated dataset was computed by switching phenotype of misclassified individuals identified as above from cases to controls.

For epilepsy and bipolar disorder datasets, ten sets of average misclassification probabilities were computed using PheLEx. For bipolar disorder, average misclassification probabilities computed showed a natural clustering and all samples with average misclassification probability > 0.5 across PheLEx runs were considered to be misclassified. For epilepsy, as estimated misclassification probabilities lacked natural clustering and were more dispersed, a more stringent criterion was applied with samples having average misclassification probability > 0.7 across at least two sets of PheLEx runs were identified as misclassified. A potential reason for dispersal could be the heterogeneous nature of epilepsy resulting in broad spread of misclassification probabilities. For both epilepsy and bipolar disorder datasets, phenotypes for samples identified as misclassified were switched from cases to controls to compute corrected phenotypes.

### Performance calculations

Performance for simulations was measured by each method’s ability to identify misclassified cases. Using misclassification probabilities computed over 100 simulated datasets at each misclassification level (5%, 10%, 20%, 30%, 40%), performance metrics such as recall/true-positive rates, false-positive rates, and precision were calculated for each method (PheLEx, PheLEx-m, PheLEx-mh and Rekaya). For decreasing threshold values in range 0.0-1.0, cases with misclassification probability higher than threshold were marked as misclassified by the method. Recall/true-positive rate was calculated as the fraction of correctly identified misclassified cases out of all misclassified cases in the dataset and false-positive rate was calculated as fraction of true cases labeled as misclassified out of all true cases. Precision was calculated as the fraction of correctly identified misclassified cases in the set of misclassified cases marked by each method. For visualization of Precision-Recall curves, mean precision across 100 simulations at each recall value was calculated per method. Similarly, for Receiver Operating Characteristic (ROC) curves mean true-positive rate across 100 simulations at each false-positive rate value was calculated per method. Area under curve for Precision-Recall curves and ROC curves were computed by calculating the area under all 100 Precision-Recall and ROC curves across misclassification levels for each method respectively.

### Measuring impact on GWAS discovery potential for simulations

To assess impact of PheLEx’s correction of phenotypes on GWAS discovery potential, recovery of disease-associated SNPs was calculated for simulations with genetic relatedness/population structure. For each simulated true phenotype, association analysis was performed before and after misclassification analysis. Disease-associated SNPs statistically significant in GWAS with simulated true phenotype (no misclassification) and misclassified phenotype were determined (according to Bonferroni-corrected p-value threshold). PheLEx was applied on simulated datasets and misclassification probabilities were computed for each phenotype. Using misclassification probabilities and methodology for identifying misclassified samples (described above), corrected phenotypes were produced for each dataset and association analysis was performed using these PheLEx corrected phenotypes to determine disease-associated SNPs that are statistically significant in GWAS with corrected phenotype. Ratios of total disease-associated SNPs significant in misclassified phenotypes versus true phenotypes (no misclassification) and those significant in corrected phenotypes versus true phenotypes (no misclassification) were computed at each misclassification rate to show recovery of disease-associated SNPs across simulations. At any misclassification rate, an increased ratio produced by corrected phenotype as compared to that produced by misclassified phenotype indicates potential of PheLEx to recover associated SNPs.

In simulations, true positive SNPs were defined as all disease-associated SNPs for the simulated true phenotype (no misclassification). True positive SNPs gained were defined as true disease-associated SNPs that were significant (according to Bonferroni-corrected p-value threshold in association analysis) in the true phenotype (no misclassification) that: (i) were not statistically significant for the misclassified phenotype, (ii) were not used as input for PheLEx, and (iii) were statistically significant in PheLEx corrected phenotypes. Novel true positive SNPs were defined as disease-associated SNPs that were not statistically significant for the true phenotype (no misclassification) or misclassified phenotype (at any misclassification rate) but were statistically significant in association analysis with PheLEx corrected phenotypes.

### Measuring impact on GWAS discovery potential for case studies

For epilepsy and bipolar disorder datasets, PheLEx was applied on these datasets to estimate misclassification probabilities and compute corrected phenotypes. P-values produced from association analysis of corrected phenotype were adjusted using the Benjamini-Hochberg procedure. SNPs that passed statistical significance threshold (adjusted p-value < 0.1) were identified and the r^2^ measure of LD was computed between these SNPs and those used for training PheLEx (training SNPs). All non-training SNPs where r^2^ < 0.1 with training SNPs and adjusted p-value < 0.1 in GWAS with the corrected phenotypes were considered ‘recovered’. Annotations for these recovered SNPs including nearest HUGO Gene Nomenclature Committee (HGNC) genes and other SNPs in LD were obtained using the web-based resource SNPsnap[98].

## Discussion

PheLEx provides two advances when considering existing methodologies[49, 73, 77] available for Bayesian misclassification analysis in GWAS: (i) PheLEx has significantly improved performance for identifying misclassified phenotypes when considering allelic effect sizes in a realistic range observed in GWAS and (ii) PheLEx provides a novel method for identifying new disease loci associations not detectable with a standard GWAS analysis. In addition, PheLEx is a complete framework that includes components that have been presented as distinct models and/or associated algorithms in the literature (i.e. differential misclassification in cases versus controls[77, 116, 117], considerations of Adaptive Metropolis-Hastings[113] and an underlying mixed model for Bayesian inference using Gibbs sampling[110, 111]). The complete PheLEx framework includes the capability to account for differential misclassification (i.e., different rates of misclassification in cases versus controls) while accounting for mixed effects due to genetic relatedness and population structure, a combination which is essential for GWAS analysis. We also provide an R package phelex to allow application of the entire PheLEx framework for GWAS analysis (see Web Resources).

When considering the application of PheLEx in extracting misclassification, there are two components of the PheLEx framework that lead to significant improvements in overall performance compared to existing methods[49, 73, 77] in datasets simulated with realistic genotype-specific odds ratios[83, 84, 103]. First, PheLEx includes an Adaptive Metropolis-Hastings step within Gibbs sampling that improves posterior sampling resulting in improved performance (Fig 2, S1 and S5 Figs). Second, the PheLEx framework uses filtered genotypes as input to filter out SNPs that have a high probability of being uninformative for learning which phenotypes are misclassified, where using more extreme p-value thresholds to prune SNPs increases the likelihood of training PheLEx on informative SNPs that will accurately point to the misclassified samples. This filtering approach not only provides dramatic savings in terms of computational expense (S6 Fig) but also improves the accuracy of misclassification estimate rates when considering realistic disease allele effect sizes (S5 and S7 Figs). Given existing methods[49, 73, 77] do not have acceptable levels of performance for misclassification analysis unless unrealistic allelic effect sizes are considered, PheLEx represents the first Bayesian misclassification that is viable for misclassification analysis for real GWAS.

The novel methodology advance in PheLEx is the use of misclassified phenotypes to re-analyze GWAS data to identify new associated loci. The joint assumptions justifying this approach are that: (i) there is at least one alternative labeling of case and control phenotypes in a GWAS (i.e., there are misclassified phenotypes) that are accurately and reproducibly reflecting the impact of alleles and, (ii) that observed genotype associations are strong enough such that there is a reasonable probability of correctly recovering these misclassified phenotypes. Clearly, these assumptions will not always be justified, such that application of this methodology should be applied judiciously (e.g., disease phenotypes known to be problematic to measure but where there are reasonably sized genotype associations) and new discoveries using this application of PheLEx should also be treated with caution. However, as demonstrated by epilepsy and bipolar disorder phenotypes, new reasonable candidate loci can be recovered with this approach. Additionally, as shown with epilepsy, this approach may also produce validation evidence for candidates identified in other studies.

Considered more broadly, the PheLEx framework is addressing a specific problem of misclassification of disease phenotypes in GWAS that is really a function of the overlapping issues of measurement error and incomplete understanding of disease etiology. These can manifest in a number of ways, including misclassification due to: (i) disease similarity and inaccurately measured current diagnosis protocols[57, 58, 118], and (ii) heterogeneous diseases or disease complexes defined as the “same” disease with a single diagnosis protocol[119-122]. The PheLEx framework we introduce is agnostic to the cause of misclassification and rather assumes that the underlying genetics can be leveraged to provide an accurate assessment of misclassification, regardless of cause, and is of value whether used purely for misclassification or for identifying novel associations. For misclassification, there is clear value in identifying healthy individuals who were misdiagnosed such that PheLEx presents an opportunity to identify “false cases” and investigate basis of their misclassification, which may relate to diagnosis error, disease subtypes or differential patterns in diagnostic measurements. Since considering a mixture of diseases with different genetics or a disease phenotype with high heterogeneity may impede extraction of disease-associated loci within a GWAS[50], PheLEx can quickly identify cases in the study that are inconsistent based on underlying disease-associated SNPs, and by defining more tractable GWAS phenotypes, boost the power of association analysis to allow users to identify new loci of interest. PheLEx therefore has promise as a novel analysis methodology for identifying candidate loci in GWAS that can be applied not only to new GWAS but also to re-analyze existing GWAS data.

## Acknowledgments

This research has been conducted using the UK Biobank Resource under Application Number 19947. We thank Andrew G. Clark and Amy L. Williams for their continued guidance and support in shaping this research.

## Supporting information

**S1 Fig. Comparison of performance between PheLEx and other methods on simulations with genetic relatedness/population structure using Receiver Operating Characteristic curves.** (A-E) Receiver Operating Characteristic (ROC) curves are shown with mean true-positive rates (y-axis) against false-positive rates (x-axis) across 100 simulations stratified with increasing misclassification rates (5%, 10%, 20%, 30% and 40%). (F) Boxplots for area under ROC curves (AUC) are shown against increasing misclassification levels across simulations for methods: (i) PheLEx-m (median AUC: 0.729, 0.748, 0.751, 0.775 and 0.785), (ii) Rekaya (median AUC: 0.596, 0.598, 0.612, 0.628 and 0.647), (iii) PheLEx (median AUC: 0.927, 0.915, 0.889, 0.856 and 0.812), and (iv) PheLEx-mh (median AUC: 0.904, 0.894, 0.856, 0.814 and 0.767). Methods are shown in different colors: PheLEx-m (dark blue), Rekaya (dark gray), PheLEx (light blue), and PheLEx-mh (light gray).

**S2 Fig. Comparison of running time between misclassification extraction methods.** Boxplots of running time (y-axis) over 500 simulation datasets with genetic relatedness/population structure (100 simulations at each of five misclassification rates) is shown for each misclassification extraction method (x-axis): (i) PheLEx-m (dark blue, median: 6.34 minutes), (ii) Rekaya (dark gray, median: 41.6 minutes), (iii) PheLEx (light blue, median: 31.2 minutes), and (iv) PheLEx-mh (light gray, median: 601 minutes).

**S3 Fig. Dataset details for simulations with random effects genetic relatedness/population structure.** (A) Histogram for Minor Allele Frequencies (MAF) (x-axis) for simulated genotypes produced using R package PhenotypeSimulator, where disease-associated genotypes are highlighted in red and gray otherwise. (B) Histogram for simulated effect sizes (x-axis) for disease-associated SNPs where effect sizes were sampled from a normal distribution with mean 2 and variance 0.3. (C) Histogram of genotype-specific odds ratio (x-axis) for simulated dataset that ranges mainly between 1.0-3.0 with a few outliers. (D) Distribution of effect sizes (y-axis) for simulated disease-associated SNPs against their minor allele frequencies (x-axis) colored using genotype-specific odds ratio value for that SNP.

**S4 Fig. Quantile-Quantile plots for p-values from GWAS for epilepsy and bipolar disorder phenotypes.** Quantile-Quantile plots (observed −log_10_ p-values (y-axis) versus expected −log_10_ p-values (x-axis)) for GWAS of (A) original epilepsy phenotype, (B) PheLEx corrected epilepsy phenotype, (C) original bipolar disorder phenotype, and (D) PheLEx corrected bipolar disorder phenotype.

**S5 Fig. Performance analysis of simulations without genetic relatedness/population structure.** (A-E) Mean Precision (y-axis) over Recall (x-axis) over 100 simulated datasets without genetic relatedness/population structure is shown across increasing misclassification (5%, 10%, 20%, 30% and 40%). Misclassification extraction methods: (i) PheLEx-m (dark blue) represents PheLEx without mixed model, (ii) Rekaya with PheLEx input (dark gray) represents methodology implemented by Rekaya et al[77] using filtration of SNPs as performed in PheLEx framework, and (iii) Rekaya (light gray) represents methodology as developed by Rekaya et al[77] with unfiltered input. (F) Boxplots for area under Precision-Recall curves (y-axis) are shown for 100 simulation datasets against increasing misclassification (x-axis) for misclassification extraction methods (Median AUC stated in order of increasing misclassification): PheLEx-m (dark blue; median AUC: 0.107, 0.195, 0.357, 0.503 and 0.585), Rekaya with PheLEx input (dark gray; median AUC: 0.0609, 0.119, 0.223, 0.322 and 0.400), and Rekaya (light gray; median AUC: 0.0701, 0.129, 0.213, 0.283 and 0.343).

**S6 Fig. Effect on performance with filtration of SNPs.** Boxplots of running time (y-axis) over simulation datasets without genetic relatedness/population structure are shown for misclassification extraction methods (x-axis): (i) PheLEx-m (dark blue, median: 7.79 minutes), (ii) Rekaya with PheLEx input (dark gray, median: 41.6 minutes) and (iii) Rekaya (light gray, median: 6504 minutes).

**S7 Fig. Dataset details for simulations without random effects (genetic relatedness/population structure).** (A) Histogram for Minor Allele Frequencies (MAF) (x-axis) for simulated genotypes used in simulation dataset produced using R package PhenotypeSimulator, where red indicates phenotype-associated genotypes and gray indicates non-phenotype associated genotypes. (B) Histogram for simulated effect sizes (x-axis) where effect sizes were sampled from a normal distribution with mean 2 and variance 0.3. (C) Histogram of genotype-specific odds ratio (x-axis) for simulated dataset that ranges mainly between 1.0-3.0 with a few outliers. (D) Distribution of effect sizes (y-axis) for simulated phenotype-associated SNPs against their minor allele frequencies (x-axis) colored using genotype-specific odds ratio value for that SNP.

**S1 Table. Distribution of attributes for UK Biobank epilepsy and bipolar disorder datasets.**

**S1 Text. Analysis of datasets simulated without genetic relatedness/population structure.**

**S2 Text. Algorithm details for PheLEx-mh, PheLEx-m and Rekaya.**

**S1 File. P-values of GWAS for original epilepsy phenotype.**

**S2 File. P-values of GWAS for PheLEx corrected epilepsy phenotype.**

**S3 File. Sample identifiers for potentially misclassified samples identified using PheLEx for UK Biobank original epilepsy phenotype.**

**S4 File. P-values of GWAS for original bipolar disorder phenotype.**

**S5 File. P-values of GWAS for PheLEx corrected bipolar disorder phenotype.**

**S6 File. Sample identifiers for potentially misclassified samples identified using PheLEx for UK Biobank bipolar disorder phenotype.**

## References

1. Price AL, Spencer CC, Donnelly P. Progress and promise in understanding the genetic basis of common diseases. Proc Biol Sci. 2015;282(1821):20151684. Epub 2015/12/25. doi:10.1098/rspb.2015.1684. PubMed PMID: 26702037; PubMed Central PMCID: PMCPMC4707742.

2. Visscher PM, Wray NR, Zhang Q, Sklar P, McCarthy MI, Brown MA, et al. 10 Years of GWAS Discovery: Biology, Function, and Translation. Am J Hum Genet. 2017;101(1):5-22. Epub 2017/07/08. doi:10.1016/j.ajhg.2017.06.005. PubMed PMID: 28686856; PubMed Central PMCID: PMCPMC5501872.

3. Schizophrenia Working Group of the Psychiatric Genomics C. Biological insights from 108 schizophrenia-associated genetic loci. Nature. 2014;511(7510):421-7. Epub 2014/07/25. doi:10.1038/nature13595. PubMed PMID: 25056061; PubMed Central PMCID: PMCPMC4112379.

4. Wellcome Trust Case Control C. Genome-wide association study of 14,000 cases of seven common diseases and 3,000 shared controls. Nature. 2007;447(7145):661-78. Epub 2007/06/08. doi:10.1038/nature05911. PubMed PMID: 17554300; PubMed Central PMCID: PMCPMC2719288.

5. Duerr RH, Taylor KD, Brant SR, Rioux JD, Silverberg MS, Daly MJ, et al. A genome-wide association study identifies IL23R as an inflammatory bowel disease gene. Science. 2006;314(5804):1461-3. Epub 2006/10/28. doi:10.1126/science.1135245. PubMed PMID: 17068223; PubMed Central PMCID: PMCPMC4410764.

6. Klein RJ, Zeiss C, Chew EY, Tsai JY, Sackler RS, Haynes C, et al. Complement factor H polymorphism in age-related macular degeneration. Science. 2005;308(5720):385-9. Epub 2005/03/12. doi:10.1126/science.1109557. PubMed PMID: 15761122; PubMed Central PMCID: PMCPMC1512523.

7. Manolio TA, Collins FS, Cox NJ, Goldstein DB, Hindorff LA, Hunter DJ, et al. Finding the missing heritability of complex diseases. Nature. 2009;461(7265):747-53. Epub 2009/10/09. doi:10.1038/nature08494. PubMed PMID: 19812666; PubMed Central PMCID: PMCPMC2831613.

8. Eichler EE, Flint J, Gibson G, Kong A, Leal SM, Moore JH, et al. Missing heritability and strategies for finding the underlying causes of complex disease. Nat Rev Genet. 2010;11(6):446-50. Epub 2010/05/19. doi:10.1038/nrg2809. PubMed PMID: 20479774; PubMed Central PMCID: PMCPMC2942068.

9. Austin MA, Hair MS, Fullerton SM. Research guidelines in the era of large-scale collaborations: an analysis of Genome-wide Association Study Consortia. Am J Epidemiol. 2012;175(9):962-9. Epub 2012/04/12. doi:10.1093/aje/kwr441. PubMed PMID: 22491085; PubMed Central PMCID: PMCPMC3339312.

10. Sullivan PF. The psychiatric GWAS consortium: big science comes to psychiatry. Neuron. 2010;68(2):182-6. Epub 2010/10/20. doi:10.1016/j.neuron.2010.10.003. PubMed PMID: 20955924; PubMed Central PMCID: PMCPMC2991765.

11. Sniekers S, Stringer S, Watanabe K, Jansen PR, Coleman JRI, Krapohl E, et al. Genome-wide association meta-analysis of 78,308 individuals identifies new loci and genes influencing human intelligence. Nat Genet. 2017;49(7):1107-12. Epub 2017/05/23. doi:10.1038/ng.3869. PubMed PMID: 28530673; PubMed Central PMCID: PMCPMC5665562.

12. Peterson RE, Edwards AC, Bacanu SA, Dick DM, Kendler KS, Webb BT. The utility of empirically assigning ancestry groups in cross-population genetic studies of addiction. Am J Addict. 2017;26(5):494-501. Epub 2017/07/18. doi:10.1111/ajad.12586. PubMed PMID: 28714599; PubMed Central PMCID: PMCPMC5646819.

13. Taylor JY, Schwander K, Kardia SL, Arnett D, Liang J, Hunt SC, et al. A Genome-wide study of blood pressure in African Americans accounting for gene-smoking interaction. Sci Rep. 2016;6:18812. Epub 2016/01/12. doi:10.1038/srep18812. PubMed PMID: 26752167; PubMed Central PMCID: PMCPMC4707536.

14. Velez Edwards DR, Naj AC, Monda K, North KE, Neuhouser M, Magvanjav O, et al. Gene-environment interactions and obesity traits among postmenopausal African-American and Hispanic women in the Women’s Health Initiative SHARe Study. Hum Genet. 2013;132(3):323-36. Epub 2012/11/30. doi:10.1007/s00439-012-1246-3. PubMed PMID: 23192594; PubMed Central PMCID: PMCPMC3704217.

15. Gao X, Nannini DR, Corrao K, Torres M, Chen YI, Fan BJ, et al. Genome-wide association study identifies WNT7B as a novel locus for central corneal thickness in Latinos. Hum Mol Genet. 2016;25(22):5035-45. Epub 2017/02/09. doi:10.1093/hmg/ddw319. PubMed PMID: 28171582; PubMed Central PMCID: PMCPMC6078592.

16. Genomes Project C, Abecasis GR, Auton A, Brooks LD, DePristo MA, Durbin RM, et al. An integrated map of genetic variation from 1,092 human genomes. Nature. 2012;491(7422):56-65. Epub 2012/11/07. doi:10.1038/nature11632. PubMed PMID: 23128226; PubMed Central PMCID: PMCPMC3498066.

17. 100,000 Genomes project. Available from: https://www.genomicsengland.co.uk/the- 100000-genomes-project/.

18. Steinthorsdottir V, Thorleifsson G, Sulem P, Helgason H, Grarup N, Sigurdsson A, et al. Identification of low-frequency and rare sequence variants associated with elevated or reduced risk of type 2 diabetes. Nat Genet. 2014;46(3):294-8. Epub 2014/01/28. doi:10.1038/ng.2882. PubMed PMID: 24464100.

19. Auer PL, Lettre G. Rare variant association studies: considerations, challenges and opportunities. Genome Med. 2015;7(1):16. Epub 2015/02/25. doi:10.1186/s13073-015-0138-2. PubMed PMID: 25709717; PubMed Central PMCID: PMCPMC4337325.

20. Long T, Hicks M, Yu HC, Biggs WH, Kirkness EF, Menni C, et al. Whole-genome sequencing identifies common-to-rare variants associated with human blood metabolites. Nat Genet. 2017;49(4):568-78. Epub 2017/03/07. doi:10.1038/ng.3809. PubMed PMID: 28263315.

21. Carss KJ, Arno G, Erwood M, Stephens J, Sanchis-Juan A, Hull S, et al. Comprehensive Rare Variant Analysis via Whole-Genome Sequencing to Determine the Molecular Pathology of Inherited Retinal Disease. Am J Hum Genet. 2017;100(1):75-90. Epub 2017/01/04. doi:10.1016/j.ajhg.2016.12.003. PubMed PMID: 28041643; PubMed Central PMCID: PMCPMC5223092.

22. Hill SR, Jr., Barker SB, Mc NJ, Tingley JO, Hibbett LL. The metabolic effects of the acetic and propionic acid analogs of thyroxine and triiodothyronine. J Clin Invest. 1960;39:523-33. Epub 1960/03/01. doi:10.1172/JCI104066. PubMed PMID: 13852407; PubMed Central PMCID: PMCPMC293333.

23. Purcell S, Neale B, Todd-Brown K, Thomas L, Ferreira MA, Bender D, et al. PLINK: a tool set for whole-genome association and population-based linkage analyses. Am J Hum Genet. 2007;81(3):559-75. doi:10.1086/519795. PubMed PMID: 17701901; PubMed Central PMCID: PMCPMC1950838.

24. Zhang Y, Liu JS. Bayesian inference of epistatic interactions in case-control studies. Nat Genet. 2007;39(9):1167-73. Epub 2007/08/28. doi:10.1038/ng2110. PubMed PMID: 17721534.

25. Winham SJ, Biernacka JM. Gene-environment interactions in genome-wide association studies: current approaches and new directions. J Child Psychol Psychiatry. 2013;54(10):1120-34. Epub 2013/07/03. doi:10.1111/jcpp.12114. PubMed PMID: 23808649; PubMed Central PMCID: PMCPMC3829379.

26. Wang T, Ho G, Ye K, Strickler H, Elston RC. A partial least-square approach for modeling gene-gene and gene-environment interactions when multiple markers are genotyped. Genet Epidemiol. 2009;33(1):6-15. Epub 2008/07/11. doi:10.1002/gepi.20351. PubMed PMID: 18615621; PubMed Central PMCID: PMCPMC2700837.

27. Tzeng JY, Zhang D, Pongpanich M, Smith C, McCarthy MI, Sale MM, et al. Studying gene and gene-environment effects of uncommon and common variants on continuous traits: a marker-set approach using gene-trait similarity regression. Am J Hum Genet. 2011;89(2):277-88. Epub 2011/08/13. doi:10.1016/j.ajhg.2011.07.007. PubMed PMID: 21835306; PubMed Central PMCID: PMCPMC3155192.

28. Thomas D. Methods for investigating gene-environment interactions in candidate pathway and genome-wide association studies. Annu Rev Public Health. 2010;31:21-36. Epub 2010/01/15. doi:10.1146/annurev.publhealth.012809.103619. PubMed PMID: 20070199; PubMed Central PMCID: PMCPMC2847610.

29. Lee S, Abecasis GR, Boehnke M, Lin X. Rare-variant association analysis: study designs and statistical tests. Am J Hum Genet. 2014;95(1):5-23. Epub 2014/07/06. doi:10.1016/j.ajhg.2014.06.009. PubMed PMID: 24995866; PubMed Central PMCID: PMCPMC4085641.

30. Li B, Leal SM. Methods for detecting associations with rare variants for common diseases: application to analysis of sequence data. Am J Hum Genet. 2008;83(3):311-21. Epub 2008/08/12. doi:10.1016/j.ajhg.2008.06.024. PubMed PMID: 18691683; PubMed Central PMCID: PMCPMC2842185.

31. Han F, Pan W. A data-adaptive sum test for disease association with multiple common or rare variants. Hum Hered. 2010;70(1):42-54. Epub 2010/04/24. doi:10.1159/000288704. PubMed PMID: 20413981; PubMed Central PMCID: PMCPMC2912645.

32. Wu MC, Lee S, Cai T, Li Y, Boehnke M, Lin X. Rare-variant association testing for sequencing data with the sequence kernel association test. Am J Hum Genet. 2011;89(1):82-93. Epub 2011/07/09. doi:10.1016/j.ajhg.2011.05.029. PubMed PMID: 21737059; PubMed Central PMCID: PMCPMC3135811.

33. Javed A, Agrawal S, Ng PC. Phen-Gen: combining phenotype and genotype to analyze rare disorders. Nat Methods. 2014;11(9):935-7. Epub 2014/08/05. doi:10.1038/nmeth.3046. PubMed PMID: 25086502.

34. International Schizophrenia C, Purcell SM, Wray NR, Stone JL, Visscher PM, O’Donovan MC, et al. Common polygenic variation contributes to risk of schizophrenia and bipolar disorder. Nature. 2009;460(7256):748-52. Epub 2009/07/03. doi:10.1038/nature08185. PubMed PMID: 19571811; PubMed Central PMCID: PMCPMC3912837.

35. Morris AP, Voight BF, Teslovich TM, Ferreira T, Segre AV, Steinthorsdottir V, et al. Large-scale association analysis provides insights into the genetic architecture and pathophysiology of type 2 diabetes. Nat Genet. 2012;44(9):981-90. Epub 2012/08/14. doi:10.1038/ng.2383. PubMed PMID: 22885922; PubMed Central PMCID: PMCPMC3442244.

36. Fuchsberger C, Flannick J, Teslovich TM, Mahajan A, Agarwala V, Gaulton KJ, et al. The genetic architecture of type 2 diabetes. Nature. 2016;536(7614):41-7. Epub 2016/07/12. doi:10.1038/nature18642. PubMed PMID: 27398621; PubMed Central PMCID: PMCPMC5034897.

37. Replication DIG, Meta-analysis C, Asian Genetic Epidemiology Network Type 2 Diabetes C, South Asian Type 2 Diabetes C, Mexican American Type 2 Diabetes C, Type 2 Diabetes Genetic Exploration by Nex-generation sequencing in muylti-Ethnic Samples C, et al. Genome-wide trans-ancestry meta-analysis provides insight into the genetic architecture of type 2 diabetes susceptibility. Nat Genet. 2014;46(3):234-44. Epub 2014/02/11. doi:10.1038/ng.2897. PubMed PMID: 24509480; PubMed Central PMCID: PMCPMC3969612.

38. van der Sluis S, Verhage M, Posthuma D, Dolan CV. Phenotypic complexity, measurement bias, and poor phenotypic resolution contribute to the missing heritability problem in genetic association studies. PLoS One. 2010;5(11):e13929. Epub 2010/11/19. doi:10.1371/journal.pone.0013929. PubMed PMID: 21085666; PubMed Central PMCID: PMCPMC2978099.

39. Cross-Disorder Phenotype Group of the Psychiatric GC, Craddock N, Kendler K, Neale M, Nurnberger J, Purcell S, et al. Dissecting the phenotype in genome-wide association studies of psychiatric illness. Br J Psychiatry. 2009;195(2):97-9. Epub 2009/08/04. doi:10.1192/bjp.bp.108.063156. PubMed PMID: 19648536; PubMed Central PMCID: PMCPMC4739810.

40. MacRae CA, Vasan RS. Next-generation genome-wide association studies: time to focus on phenotype? Circ Cardiovasc Genet. 2011;4(4):334-6. Epub 2011/08/19. doi:10.1161/CIRCGENETICS.111.960765. PubMed PMID: 21846867; PubMed Central PMCID: PMCPMC3187849.

41. Gage JL, de Leon N, Clayton MK. Comparing Genome-Wide Association Study Results from Different Measurements of an Underlying Phenotype. G3 (Bethesda). 2018;8(11):3715-22. Epub 2018/09/29. doi:10.1534/g3.118.200700. PubMed PMID: 30262522; PubMed Central PMCID: PMCPMC6222562.

42. Ronnegard L, McFarlane SE, Husby A, Kawakami T, Ellegren H, Qvarnstrom A. Increasing the power of genome wide association studies in natural populations using repeated measures - evaluation and implementation. Methods Ecol Evol. 2016;7(7):792-9. Epub 2016/08/02. doi:10.1111/2041-210X.12535. PubMed PMID: 27478587; PubMed Central PMCID: PMCPMC4950150.

43. Barendse W. The effect of measurement error of phenotypes on genome wide association studies. BMC Genomics. 2011;12:232. Epub 2011/05/17. doi:10.1186/1471-2164-12-232. PubMed PMID: 21569388; PubMed Central PMCID: PMCPMC3224145.

44. Maier RM, Zhu Z, Lee SH, Trzaskowski M, Ruderfer DM, Stahl EA, et al. Improving genetic prediction by leveraging genetic correlations among human diseases and traits. Nat Commun. 2018;9(1):989. Epub 2018/03/09. doi:10.1038/s41467-017-02769-6. PubMed PMID: 29515099; PubMed Central PMCID: PMCPMC5841449.

45. Schifano ED, Li L, Christiani DC, Lin X. Genome-wide association analysis for multiple continuous secondary phenotypes. Am J Hum Genet. 2013;92(5):744-59. Epub 2013/05/07. doi:10.1016/j.ajhg.2013.04.004. PubMed PMID: 23643383; PubMed Central PMCID: PMCPMC3644646.

46. Fusi N, Lippert C, Lawrence ND, Stegle O. Warped linear mixed models for the genetic analysis of transformed phenotypes. Nat Commun. 2014;5:4890. Epub 2014/09/23. doi:10.1038/ncomms5890. PubMed PMID: 25234577; PubMed Central PMCID: PMCPMC4199105.

47. Valenstein PN. Evaluating diagnostic tests with imperfect standards. Am J Clin Pathol. 1990;93(2):252-8. Epub 1990/02/01. PubMed PMID: 2405632.

48. Rutjes AW, Reitsma JB, Coomarasamy A, Khan KS, Bossuyt PM. Evaluation of diagnostic tests when there is no gold standard. A review of methods. Health Technol Assess. 2007;11(50):iii, ix-51. Epub 2007/11/21. PubMed PMID: 18021577.

49. Smith S, Hay el H, Farhat N, Rekaya R. Genome wide association studies in presence of misclassified binary responses. BMC Genet. 2013;14:124. doi:10.1186/1471-2156-14-124. PubMed PMID: 24369108; PubMed Central PMCID: PMCPMC3879434.

50. Manchia M, Cullis J, Turecki G, Rouleau GA, Uher R, Alda M. The impact of phenotypic and genetic heterogeneity on results of genome wide association studies of complex diseases. PLoS One. 2013;8(10):e76295. Epub 2013/10/23. doi:10.1371/journal.pone.0076295. PubMed PMID: 24146854; PubMed Central PMCID: PMCPMC3795757.

51. Gordon D, Yang Y, Haynes C, Finch SJ, Mendell NR, Brown AM, et al. Increasing power for tests of genetic association in the presence of phenotype and/or genotype error by use of double-sampling. Stat Appl Genet Mol Biol. 2004;3:Article26. Epub 2006/05/02. doi:10.2202/1544-6115.1085. PubMed PMID: 16646805.

52. Edwards BJ, Haynes C, Levenstien MA, Finch SJ, Gordon D. Power and sample size calculations in the presence of phenotype errors for case/control genetic association studies. BMC Genet. 2005;6:18. Epub 2005/04/12. doi:10.1186/1471-2156-6-18. PubMed PMID: 15819990; PubMed Central PMCID: PMCPMC1131899.

53. Ji F, Yang Y, Haynes C, Finch SJ, Gordon D. Computing asymptotic power and sample size for case-control genetic association studies in the presence of phenotype and/or genotype misclassification errors. Stat Appl Genet Mol Biol. 2005;4:Article37. Epub 2006/05/02. doi:10.2202/1544-6115.1184. PubMed PMID: 16646856.

54. Barral S, Haynes C, Stone M, Gordon D. LRTae: improving statistical power for genetic association with case/control data when phenotype and/or genotype misclassification errors are present. BMC Genet. 2006;7:24. Epub 2006/05/13. doi:10.1186/1471-2156-7-24. PubMed PMID: 16689984; PubMed Central PMCID: PMCPMC1471798.

55. Gordon D, Haynes C, Yang Y, Kramer PL, Finch SJ. Linear trend tests for case-control genetic association that incorporate random phenotype and genotype misclassification error. Genet Epidemiol. 2007;31(8):853-70. Epub 2007/06/15. doi:10.1002/gepi.20246. PubMed PMID: 17565750.

56. Buyske S, Yang G, Matise TC, Gordon D. When a case is not a case: effects of phenotype misclassification on power and sample size requirements for the transmission disequilibrium test with affected child trios. Hum Hered. 2009;67(4):287-92. Epub 2009/01/28. doi:10.1159/000194981. PubMed PMID: 19172087.

57. Winnie Qian WQ, Tom Schweizer, David Munoz, Corinne E. Fischer. Misdiagnosis Of Alzheimer’s Disease: Inconsistencies Between Clinical Diagnosis And Neuropathological Confirmation. Elsevier. 2016;12(7):P293.

58. Gaugler JE, Ascher-Svanum H, Roth DL, Fafowora T, Siderowf A, Beach TG. Characteristics of patients misdiagnosed with Alzheimer’s disease and their medication use: an analysis of the NACC-UDS database. BMC Geriatr. 2013;13:137. Epub 2013/12/21. doi:10.1186/1471-2318-13-137. PubMed PMID: 24354549; PubMed Central PMCID: PMCPMC3878261.

59. Bromet EJ, Kotov R, Fochtmann LJ, Carlson GA, Tanenberg-Karant M, Ruggero C, et al. Diagnostic shifts during the decade following first admission for psychosis. Am J Psychiatry. 2011;168(11):1186-94. doi:10.1176/appi.ajp.2011.11010048. PubMed PMID: 21676994; PubMed Central PMCID: PMCPMC3589618.

60. Singh T, Rajput M. Misdiagnosis of bipolar disorder. Psychiatry (Edgmont). 2006;3(10):57-63. PubMed PMID: 20877548; PubMed Central PMCID: PMCPMC2945875.

61. Ghaemi SN, Sachs GS, Chiou AM, Pandurangi AK, Goodwin K. Is bipolar disorder still underdiagnosed? Are antidepressants overutilized? J Affect Disord. 1999;52(1-3):135-44. PubMed PMID: 10357026.

62. Ghaemi SN, Boiman EE, Goodwin FK. Diagnosing bipolar disorder and the effect of antidepressants: a naturalistic study. J Clin Psychiatry. 2000;61(10):804-8; quiz 9. PubMed PMID: 11078046.

63. Hirschfeld RM, Lewis L, Vornik LA. Perceptions and impact of bipolar disorder: how far have we really come? Results of the national depressive and manic-depressive association 2000 survey of individuals with bipolar disorder. J Clin Psychiatry. 2003;64(2):161-74. Epub 2003/03/14. PubMed PMID: 12633125.

64. Ghouse AA, Sanches M, Zunta-Soares G, Swann AC, Soares JC. Overdiagnosis of bipolar disorder: a critical analysis of the literature. ScientificWorldJournal. 2013;2013:297087. Epub 2013/12/19. doi:10.1155/2013/297087. PubMed PMID: 24348150; PubMed Central PMCID: PMCPMC3856145.

65. Solomon AJ, Bourdette DN, Cross AH, Applebee A, Skidd PM, Howard DB, et al. The contemporary spectrum of multiple sclerosis misdiagnosis: A multicenter study. Neurology. 2016;87(13):1393-9. Epub 2016/09/02. doi:10.1212/WNL.0000000000003152. PubMed PMID: 27581217; PubMed Central PMCID: PMCPMC5047038.

66. O’Reilly PF, Hoggart CJ, Pomyen Y, Calboli FC, Elliott P, Jarvelin MR, et al. MultiPhen: joint model of multiple phenotypes can increase discovery in GWAS. PLoS One. 2012;7(5):e34861. Epub 2012/05/09. doi:10.1371/journal.pone.0034861. PubMed PMID: 22567092; PubMed Central PMCID: PMCPMC3342314.

67. Panoutsopoulou K, Thiagarajah S, Zengini E, Day-Williams AG, Ramos YF, Meessen JM, et al. Radiographic endophenotyping in hip osteoarthritis improves the precision of genetic association analysis. Ann Rheum Dis. 2017;76(7):1199-206. Epub 2016/12/16. doi:10.1136/annrheumdis-2016-210373. PubMed PMID: 27974301; PubMed Central PMCID: PMCPMC5530347.

68. Warde-Farley D, Brudno M, Morris Q, Goldenberg A. Mixture model for sub-phenotyping in GWAS. Pac Symp Biocomput. 2012:363-74. Epub 2011/12/17. PubMed PMID: 22174291.

69. Yang JJ, Williams LK, Buu A. Identifying Pleiotropic Genes in Genome-Wide Association Studies for Multivariate Phenotypes with Mixed Measurement Scales. PLoS One. 2017;12(1):e0169893. Epub 2017/01/13. doi:10.1371/journal.pone.0169893. PubMed PMID: 28081206; PubMed Central PMCID: PMCPMC5231271.

70. Duffy SW, Warwick J, Williams AR, Keshavarz H, Kaffashian F, Rohan TE, et al. A simple model for potential use with a misclassified binary outcome in epidemiology. J Epidemiol Community Health. 2004;58(8):712-7. Epub 2004/07/15. doi:10.1136/jech.2003.010546. PubMed PMID: 15252078; PubMed Central PMCID: PMCPMC1732865.

71. Prescott GJ, Garthwaite PH. A Bayesian approach to prospective binary outcome studies with misclassification in a binary risk factor. Stat Med. 2005;24(22):3463-77. Epub 2005/10/21. doi:10.1002/sim.2192. PubMed PMID: 16237661.

72. Magder LS, Hughes JP. Logistic regression when the outcome is measured with uncertainty. Am J Epidemiol. 1997;146(2):195-203. Epub 1997/07/15. PubMed PMID: 9230782.

73. Rekaya R, Smith S, Hay el H, Aggrey SE. Misclassification in binary responses and effect on genome-wide association studies. Poult Sci. 2013;92(9):2535-40. Epub 2013/08/21. doi:10.3382/ps.2012-02738. PubMed PMID: 23960139.

74. Hofler M. The effect of misclassification on the estimation of association: a review. Int J Methods Psychiatr Res. 2005;14(2):92-101. Epub 2005/09/24. PubMed PMID: 16175878.

75. Graber ML. The incidence of diagnostic error in medicine. BMJ Qual Saf. 2013;22 Suppl 2:ii21-ii7. Epub 2013/06/19. doi:10.1136/bmjqs-2012-001615. PubMed PMID: 23771902; PubMed Central PMCID: PMCPMC3786666.

76. Singh H, Schiff GD, Graber ML, Onakpoya I, Thompson MJ. The global burden of diagnostic errors in primary care. BMJ Qual Saf. 2017;26(6):484-94. Epub 2016/08/18. doi:10.1136/bmjqs-2016-005401. PubMed PMID: 27530239; PubMed Central PMCID: PMCPMC5502242.

77. Rekaya R, Smith S, Hay EH, Farhat N, Aggrey SE. Analysis of binary responses with outcome-specific misclassification probability in genome-wide association studies. Appl Clin Genet. 2016;9:169-77. doi:10.2147/TACG.S122250. PubMed PMID: 27942229; PubMed Central PMCID: PMCPMC5138056.

78. Joseph S, Robbins K, Zhang W, Rekaya R. Effects of misdiagnosis in input data on the identification of differential expression genes in incipient Alzheimer patients. In Silico Biol. 2008;8(5-6):545-54. PubMed PMID: 19374137.

79. Joseph S, Robbins KR, Rekaya R. A statistical and biological approach for identifying misdiagnosis of incipient Alzheimer patients using gene expression data. Conf Proc IEEE Eng Med Biol Soc. 2006;1:5854-7. Epub 2007/10/20. doi:10.1109/IEMBS.2006.259371. PubMed PMID: 17947171.

80. Zhang W, Rekaya R, Bertrand K. A method for predicting disease subtypes in presence of misclassification among training samples using gene expression: application to human breast cancer. Bioinformatics. 2006;22(3):317-25. doi:10.1093/bioinformatics/bti738. PubMed PMID: 16267079.

81. Kang HM, Sul JH, Service SK, Zaitlen NA, Kong SY, Freimer NB, et al. Variance component model to account for sample structure in genome-wide association studies. Nat Genet. 2010;42(4):348-54. doi:10.1038/ng.548. PubMed PMID: 20208533; PubMed Central PMCID: PMCPMC3092069.

82. Newman DL, Abney M, McPeek MS, Ober C, Cox NJ. The importance of genealogy in determining genetic associations with complex traits. Am J Hum Genet. 2001;69(5):1146-8. Epub 2001/10/09. doi:10.1086/323659. PubMed PMID: 11590549; PubMed Central PMCID: PMCPMC1274359.

83. Luedeke M, Coinac I, Linnert CM, Bogdanova N, Rinckleb AE, Schrader M, et al. Prostate cancer risk is not altered by TP53AIP1 germline mutations in a German case-control series. PLoS One. 2012;7(3):e34128. Epub 2012/03/30. doi:10.1371/journal.pone.0034128. PubMed PMID: 22457820; PubMed Central PMCID: PMCPMC3311578.

84. Helgadottir A, Thorleifsson G, Magnusson KP, Gretarsdottir S, Steinthorsdottir V, Manolescu A, et al. The same sequence variant on 9p21 associates with myocardial infarction, abdominal aortic aneurysm and intracranial aneurysm. Nat Genet. 2008;40(2):217-24. Epub 2008/01/08. doi:10.1038/ng.72. PubMed PMID: 18176561.

85. Jakobsdottir J, Gorin MB, Conley YP, Ferrell RE, Weeks DE. Interpretation of genetic association studies: markers with replicated highly significant odds ratios may be poor classifiers. PLoS Genet. 2009;5(2):e1000337. Epub 2009/02/07. doi:10.1371/journal.pgen.1000337. PubMed PMID: 19197355; PubMed Central PMCID: PMCPMC2629574 Pittsburgh for the LOC387715/ARMS2 locus.

86. Helgason A, Yngvadottir B, Hrafnkelsson B, Gulcher J, Stefansson K. An Icelandic example of the impact of population structure on association studies. Nat Genet. 2005;37(1):90-5. Epub 2004/12/21. doi:10.1038/ng1492. PubMed PMID: 15608637.

87. Canela-Xandri O, Rawlik K, Tenesa A. An atlas of genetic associations in UK Biobank. Nat Genet. 2018;50(11):1593-9. Epub 2018/10/24. doi:10.1038/s41588-018-0248-z. PubMed PMID: 30349118.

88. Potkin SG, Guffanti G, Lakatos A, Turner JA, Kruggel F, Fallon JH, et al. Hippocampal atrophy as a quantitative trait in a genome-wide association study identifying novel susceptibility genes for Alzheimer’s disease. PLoS One. 2009;4(8):e6501. Epub 2009/08/12. doi:10.1371/journal.pone.0006501. PubMed PMID: 19668339; PubMed Central PMCID: PMCPMC2719581.

89. Bonilha L, Halford JJ, Morgan PS, Edwards JC. Hippocampal atrophy in temporal lobe epilepsy: the ‘generator’ and ‘receiver’. Acta Neurol Scand. 2012;125(2):105-10. Epub 2011/04/08. doi:10.1111/j.1600-0404.2011.01510.x. PubMed PMID: 21470191.

90. Kanner AM. Hippocampal atrophy: another common pathogenic mechanism of depressive disorders and epilepsy? Epilepsy Curr. 2011;11(5):149-50. Epub 2011/10/25. doi:10.5698/1535-7511-11.5.149. PubMed PMID: 22020737; PubMed Central PMCID: PMCPMC3193098.

91. Boudry-Labis E, Demeer B, Le Caignec C, Isidor B, Mathieu-Dramard M, Plessis G, et al. A novel microdeletion syndrome at 9q21.13 characterised by mental retardation, speech delay, epilepsy and characteristic facial features. Eur J Med Genet. 2013;56(3):163-70. Epub 2013/01/03. doi:10.1016/j.ejmg.2012.12.006. PubMed PMID: 23279911.

92. Wang J, Lin ZJ, Liu L, Xu HQ, Shi YW, Yi YH, et al. Epilepsy-associated genes. Seizure. 2017;44:11-20. Epub 2016/12/23. doi:10.1016/j.seizure.2016.11.030. PubMed PMID: 28007376.

93. Bergren SK, Rutter ED, Kearney JA. Fine mapping of an epilepsy modifier gene on mouse Chromosome 19. Mamm Genome. 2009;20(6):359-66. Epub 2009/06/11. doi:10.1007/s00335-009-9193-6. PubMed PMID: 19513789; PubMed Central PMCID: PMCPMC2804848.

94. Liu Z, Li Z, Zhi X, Du Y, Lin Z, Wu J. Identification of De Novo DNMT3A Mutations That Cause West Syndrome by Using Whole-Exome Sequencing. Mol Neurobiol. 2018;55(3):2483-93. Epub 2017/04/08. doi:10.1007/s12035-017-0483-9. PubMed PMID: 28386848.

95. Baglietto MG, Caridi G, Gimelli G, Mancardi M, Prato G, Ronchetto P, et al. RORB gene and 9q21.13 microdeletion: report on a patient with epilepsy and mild intellectual disability. Eur J Med Genet. 2014;57(1):44-6. Epub 2013/12/21. doi:10.1016/j.ejmg.2013.12.001. PubMed PMID: 24355400.

96. Reif A, Nguyen TT, Weissflog L, Jacob CP, Romanos M, Renner TJ, et al. DIRAS2 is associated with adult ADHD, related traits, and co-morbid disorders. Neuropsychopharmacology. 2011;36(11):2318-27. Epub 2011/07/14. doi:10.1038/npp.2011.120. PubMed PMID: 21750579; PubMed Central PMCID: PMCPMC3176568.

97. Torres CM, Siebert M, Bock H, Mota SM, Krammer BR, Duarte JA, et al. NTRK2 (TrkB gene) variants and temporal lobe epilepsy: A genetic association study. Epilepsy Res. 2017;137:1-8. Epub 2017/09/02. doi:10.1016/j.eplepsyres.2017.08.010. PubMed PMID: 28863320.

98. Pers TH, Timshel P, Hirschhorn JN. SNPsnap: a Web-based tool for identification and annotation of matched SNPs. Bioinformatics. 2015;31(3):418-20. Epub 2014/10/16. doi:10.1093/bioinformatics/btu655. PubMed PMID: 25316677; PubMed Central PMCID: PMCPMC4308663.

99. Perroud N, Dayer A, Piguet C, Nallet A, Favre S, Malafosse A, et al. Childhood maltreatment and methylation of the glucocorticoid receptor gene NR3C1 in bipolar disorder. Br J Psychiatry. 2014;204(1):30-5. Epub 2013/06/08. doi:10.1192/bjp.bp.112.120055. PubMed PMID: 23743517.

100. Gibbons A, Bell L, Udawela M, Dean B. mRNA expression of the P5 ATPase ATP13A4 is increased in Broca’s Area from subjects with schizophrenia. World J Biol Psychiatry. 2018:1-23. Epub 2018/12/07. doi:10.1080/15622975.2018.1548781. PubMed PMID: 30501451.

101. Scarr E, Udawela M, Thomas EA, Dean B. Changed gene expression in subjects with schizophrenia and low cortical muscarinic M1 receptors predicts disrupted upstream pathways interacting with that receptor. Mol Psychiatry. 2018;23(2):295-303. Epub 2016/11/02. doi:10.1038/mp.2016.195. PubMed PMID: 27801890; PubMed Central PMCID: PMCPMC5794886.

102. Askland K, Read C, O’Connell C, Moore JH. Ion channels and schizophrenia: a gene set-based analytic approach to GWAS data for biological hypothesis testing. Hum Genet. 2012;131(3):373-91. Epub 2011/08/26. doi:10.1007/s00439-011-1082-x. PubMed PMID: 21866342; PubMed Central PMCID: PMCPMC3278516.

103. Olsen L, Hansen T, Jakobsen KD, Djurovic S, Melle I, Agartz I, et al. The estrogen hypothesis of schizophrenia implicates glucose metabolism: association study in three independent samples. BMC Med Genet. 2008;9:39. Epub 2008/05/08. doi:10.1186/1471-2350-9-39. PubMed PMID: 18460190; PubMed Central PMCID: PMCPMC2391158.

104. Breen MS, Uhlmann A, Nday CM, Glatt SJ, Mitt M, Metsalpu A, et al. Candidate gene networks and blood biomarkers of methamphetamine-associated psychosis: an integrative RNA-sequencing report. Transl Psychiatry. 2016;6:e802. Epub 2016/05/11. doi:10.1038/tp.2016.67. PubMed PMID: 27163203; PubMed Central PMCID: PMCPMC5070070.

105. Meyer HV, Birney E. PhenotypeSimulator: A comprehensive framework for simulating multi-trait, multi-locus genotype to phenotype relationships. Bioinformatics. 2018;34(17):2951-6. Epub 2018/04/05. doi:10.1093/bioinformatics/bty197. PubMed PMID: 29617944; PubMed Central PMCID: PMCPMC6129313.

106. Chen W, Chen X, Archer KJ, Liu N, Li Q, Zhao Z, et al. A rapid association test procedure robust under different genetic models accounting for population stratification. Hum Hered. 2013;75(1):23-33. Epub 2013/04/11. doi:10.1159/000350109. PubMed PMID: 23571404; PubMed Central PMCID: PMCPMC3786013.

107. Pei YF, Zhang L, Papasian CJ, Wang YP, Deng HW. On individual genome-wide association studies and their meta-analysis. Hum Genet. 2014;133(3):265-79. Epub 2013/10/12. doi:10.1007/s00439-013-1366-4. PubMed PMID: 24114349; PubMed Central PMCID: PMCPMC4127980.

108. Mugo JW, Geza E, Defo J, Elsheikh SSM, Mazandu GK, Mulder NJ, et al. A multi-scenario genome-wide medical population genetics simulation framework. Bioinformatics. 2017;33(19):2995-3002. Epub 2017/09/29. doi:10.1093/bioinformatics/btx369. PubMed PMID: 28957497; PubMed Central PMCID: PMCPMC5870573.

109. Sudlow C, Gallacher J, Allen N, Beral V, Burton P, Danesh J, et al. UK biobank: an open access resource for identifying the causes of a wide range of complex diseases of middle and old age. PLoS Med. 2015;12(3):e1001779. doi:10.1371/journal.pmed.1001779. PubMed PMID: 25826379; PubMed Central PMCID: PMCPMC4380465.

110. Sorensen DA, Andersen, S., Gianola, D., and Korsgaard, I. Bayesian inference in threshold models using Gibbs sampling. Genetics, Selection, Evolution. 1995;17:229–49.

111. CS Wang JR, and D Gianola. Bayesian analysis of mixed linear models via Gibbs sampling with an application to litter size in Iberian pigs. Genetics Selection Evolution. 1994;26(2):91–115.

112. Tier IHaB. Estimation of variance components of threshold characters by marginal posterior modes and means via Gibbs sampling. Genetics Selection Evolution. 1995;27:519.

113. Brooks S. GA, Jones G., and Meng X.-L. MCMC Handbook 2010.

114. Gelman A. GWR, and Roberts G. O. Weak convergence and optimal scaling of random walk Metropolis algorithms. Ann Appl Probab. 1997;7(1):110–20.

115. Hoffman GE, Mezey JG, Schadt EE. lrgpr: interactive linear mixed model analysis of genome-wide association studies with composite hypothesis testing and regression diagnostics in R. Bioinformatics. 2014;30(21):3134-5. doi:10.1093/bioinformatics/btu435. PubMed PMID: 25035399; PubMed Central PMCID: PMCPMC4201153.

116. Gilbert R, Martin RM, Donovan J, Lane JA, Hamdy F, Neal DE, et al. Misclassification of outcome in case-control studies: Methods for sensitivity analysis. Stat Methods Med Res. 2016;25(5):2377-93. Epub 2014/09/14. doi:10.1177/0962280214523192. PubMed PMID: 25217446.

117. De Smedt T, Merrall E, Macina D, Perez-Vilar S, Andrews N, Bollaerts K. Bias due to differential and non-differential disease- and exposure misclassification in studies of vaccine effectiveness. PLoS One. 2018;13(6):e0199180. Epub 2018/06/16. doi:10.1371/journal.pone.0199180. PubMed PMID: 29906276; PubMed Central PMCID: PMCPMC6003693 Both companies develop vaccines and support the IMI ADVANCE project. KB received consulting fees from vaccine producing companies (GSK, SPMSD, Pfizer, Takeda) not related to the research presented here. TDS received consulting fees from Pfizer and Takeda not related to this work. At the time of the research, SPV was employed by FISABIO and Erasmus MC. The authors have declared that no competing interests exist.

118. Wang L, Chen H, Shi J, Tang H, Li H, Zheng W, et al. Castleman disease mimicking systemic lupus erythematosus: A case report. Medicine (Baltimore). 2018;97(38):e12291. Epub 2018/09/22. doi:10.1097/MD.0000000000012291. PubMed PMID: 30235674; PubMed Central PMCID: PMCPMC6160051.

119. Manson JJ, Rahman A. Systemic lupus erythematosus. Orphanet J Rare Dis. 2006;1:6. Epub 2006/05/26. doi:10.1186/1750-1172-1-6. PubMed PMID: 16722594; PubMed Central PMCID: PMCPMC1459118.

120. Wang Z, Chang C, Peng M, Lu Q. Translating epigenetics into clinic: focus on lupus. Clin Epigenetics. 2017;9:78. Epub 2017/08/09. doi:10.1186/s13148-017-0378-7. PubMed PMID: 28785369; PubMed Central PMCID: PMCPMC5541721.

121. Au R, Piers RJ, Lancashire L. Back to the future: Alzheimer’s disease heterogeneity revisited. Alzheimers Dement (Amst). 2015;1(3):368-70. Epub 2016/04/15. doi:10.1016/j.dadm.2015.05.006. PubMed PMID: 27077132; PubMed Central PMCID: PMCPMC4827150.

122. Lam B, Masellis M, Freedman M, Stuss DT, Black SE. Clinical, imaging, and pathological heterogeneity of the Alzheimer’s disease syndrome. Alzheimers Res Ther. 2013;5(1):1. Epub 2013/01/11. doi:10.1186/alzrt155. PubMed PMID: 23302773; PubMed Central PMCID: PMCPMC3580331.

